# Limb contraction drives fear perception

**DOI:** 10.1101/2020.01.09.899849

**Authors:** Marta Poyo Solanas, Maarten Vaessen, Beatrice de Gelder

**Author notes:** Address correspondence to Beatrice de Gelder, Department of Cognitive Neuroscience, Brain and Emotion Laboratory, Faculty of Psychology and Neuroscience, Maastricht University, Oxfordlaan 55, 6229 EV Maastricht, The Netherlands.

## Abstract

Humans and other primate species are experts at recognizing affective information from body movements but the underlying brain mechanisms are still largely unknown. Previous research focusing on the brain representation of symbolic emotion categories has led to mixed results. This study used representational similarity and multi-voxel pattern analysis techniques to investigate how postural and kinematic features computed from affective whole-body movement videos are related to brain processes. We show that body posture and kinematics differentially activated brain regions indicating that this information might be selectively encoded in these regions. Most specifically, the feature limb contraction seemed to be particularly relevant for distinguishing fear and it was represented in several regions spanning affective, action observation and motor preparation networks. Our approach goes beyond traditional methods of mapping symbolic emotion categories to brain activation/deactivation by discovering which specific movement features are encoded in the brain, and possibly drive automatic emotion perception.

## Introduction

It is widely agreed that humans and other primate species are experts at recognizing information from body movements. Presumably, our brains transform this visual information into an understanding of the intent and the emotion expressed (de Gelder, 2006; Giese & Rizzolatti, 2015). The central importance of nonverbal communication across many social species suggests that the brain is equipped for rapid and accurate body movement recognition; yet, the mechanisms underlying this ability are still largely unclear. Previous research has predominantly investigated the brain correlates of symbolic emotion categories and led to mixed results (Kirby & Robinson, 2017; Lindquist, Wager, Kober, Bliss-Moreau, & Barrett, 2012). A better understanding of the processes underlying the perception of affective body movements requires understanding what are the exact visual properties that drive movement and emotion perception (e.g. kinematic and postural body features) and their relation to the different brain regions involved in perception. Understanding affective body perception at the subsymbolic level can now take advantage of novel computational methods and techniques for stimulus analysis as well as for brain activity decoding. However, an important difficulty for uncovering relationships between features of body movements and the emotions expressed is the fact that the moving human body represents a complex, high-dimensional visual stimulus (Roether, Omlor, Christensen, & Giese, 2009). This raises the question of whether it is possible to extract systematically a limited set of highly informative features upon which recognition and understanding are based.

So far, there is neither a common theoretical perspective on whole-body movement analysis nor there has been an effort to integrate neural and computational findings (Kleinsmith & Bianchi-Berthouze, 2012). In the field of Neuroscience, research has mainly focused on identifying body-, emotion- and action-related areas, but it is not yet known which information is actually processed in these areas or how it contributes to emotional recognition. At the other end is Computer Vision, where the automatic recognition of body movements is of great interest. Research in this field has provided insights about postural and kinematic features important for the automatic recognition of actions and emotions expressed by the body, but without integrating evolutionary and ecological constraints of brain function.

Past studies have provided some indications about relevant features and their relation to emotional expressions. For example, it has been shown that particular body postures and movements are characteristic of distinct emotions (De Meijer, 1989; Kleinsmith & Bianchi-Berthouze, 2012; Patwardhan, 2017; Piana, Stagliano, Odone, Verri, & Camurri, 2014; Roether et al., 2009; Wallbott, 1998). In studies using static body postures as well as affective body movements from dance and gait, some important postural features have been identified, including elbow flexion, associated with the expression of anger, or head inclination, typically observed for sadness (Coulson, 2004; Vaessen, Abassi, Mancini, Camurri, & de Gelder, 2018; Wallbott, 1998). Other form-related cues that have been investigated are the vertical extension of the body (e.g. upper limbs remain low for sadness but high for happiness), the directionality of the movement (e.g. angry bodies are usually accompanied by a forward movement), symmetry (e.g. the movement of the upper limbs tends to be symmetrical when experiencing joy) or the amount of lateral opening of the body (e.g. hands are close to the body during fear and sadness while extended in happiness) (for a review see Kleinsmith & Bianchi-Berthouze, 2012). The use of point-light displays, where the only form information is derived from motion-mediated structural cues (Johansson, 1973), has been crucial in revealing the strong impact that movement kinematics, such as velocity and acceleration, have on the recognition of emotion from arm (Paterson, Pollick, & Sanford, 2001; Pollick, Paterson, Bruderlin, & Sanford, 2001; Sawada, Suda, & Ishii, 2003) and whole-body movements (Roether et al., 2009).

Although various candidate features have been studied, there is currently no clear understanding of which features are critical for conveying emotions with whole-body movements. We are equally still in the dark in understanding the relation between candidate features and brain processes. In an effort to relate questions of body movement perception to research on movement perception in general, it was suggested that two major sources of affective information, form and movement, are processed in two separate pathways in the brain corresponding to the two-stream model of visual processing (Giese & Poggio, 2003; Milner & Goodale, 2006, 2008; Vaina, Lemay, Bienfang, Choi, & Nakayama, 1990). From the primary visual cortex, the dorsal stream leads to the parietal lobe and is specialized in localizing objects in space, processing motion signals and in visual-spatial guidance of actions. The ventral stream leads to the temporal lobe and is responsible for visual form processing and object recognition. Two areas in this pathway have been identified that sustain a certain level of specialization in the processing of whole bodies and body parts: the extrastriate body area (EBA) in the medial occipital cortex, and the fusiform body area (FBA) in the fusiform gyrus (Downing, Jiang, Shuman, & Kanwisher, 2001; Peelen & Downing, 2005; Schwarzlose, Baker, & Kanwisher, 2005). However, the perception and recognition of human bodies elicits a widespread neural response beyond the visual analysis of body features (de Gelder, 2006; Van den Stock et al., 2011). Body form and movement convey a wide range of information, such emotion, action and intention. Thus, their perception also triggers the activation of areas responsible for the processing of their affective content, the conveyed action and for the preparation of an appropriate behavioural response (de Gelder, Snyder, Greve, Gerard, & Hadjikhani, 2004; Van den Stock et al., 2011).

Indeed, the observation of body postures and movements strongly activates the so-called ‘action observation network’ (AON). This network has been suggested to play an important role in the understanding of other people’s actions and their underlying intentions (Fogassi et al., 2005; Iacoboni et al., 2005; Jellema, Baker, Wicker, & Perrett, 2000; Rizzolatti, Fogassi, & Gallese, 2001). It is comprised of the ventral premotor/caudal inferior frontal gyrus complex (PMv/cIFG), the supplementary motor area (SMA), the posterior superior temporal sulcus (pSTS) and the inferior parietal lobule (IPL) (Buccino et al., 2001; Caspers, Zilles, Laird, & Eickhoff, 2010; Decety et al., 1997; Grafton, Arbib, Fadiga, & Rizzolatti, 1996). A recent body of work has also involved this network in the processing of emotional body expressions. Specifically, higher activity has been reported in several nodes of the AON for emotional body actions as opposed to neutral ones (Grèzes, Pichon, & de Gelder, 2007; Kret, Pichon, Grèzes, & de Gelder, 2011a, 2011b; Pichon, de Gelder, & Grèzes, 2009). Recent Transcranial Magnetic Stimulation (TMS) studies have directly implicated both the IPL (Engelen, de Graaf, Sack, & de Gelder, 2015) and the pSTS (Candidi, Stienen, Aglioti, & de Gelder, 2011) in the recognition of fearful body expressions. Interestingly, the pSTS also appears to be involved in processing body motion cues (Grossman, Jardine, & Pyles, 2010).

In addition to the AON, ALE meta-analyses have shown that the observation of emotional expressions involves the activity of other areas including the dorsal and ventral medial prefrontal cortices, orbitofrontal cortex, anterior cingulate cortex (ACC), posterior cingulate, insula, pre-supplementary motor area (pre-SMA) and temporal pole, and also of subcortical areas including ventral striatum, amygdala, thalamus and hypothalamus (Dricu & Frühholz, 2016; Kober et al., 2008).

However, it remains unclear whether areas selective for body movements represent affective content by encoding body form and kinematic properties. To investigate this question, this study used video clips of whole-body movements expressing anger, happiness, fear or a non-emotional action. Each actor’s joint positions were estimated using the state-of-the-art 2D pose estimation library OpenPose (Cao, Simon, Wei, & Sheikh, 2017). Quantitative features were derived from the joint 2D positions such as velocity, acceleration, vertical movement, the angles between limbs, symmetry, limb contraction and surface, giving their importance in previous work (for a review see Kleinsmith & Bianchi-Berthouze, 2012). By means of representational similarity multi-voxel pattern analysis techniques, we investigated whether the (dis)similarity of body posture and kinematics between different emotional categories could explain the neural response in and beyond body-selective regions.

## Materials and methods

### Participants

Thirteen healthy participants (mean age = 25.8; age range = 21-30; three males) took part in the experiment. All participants had normal or corrected-to-normal vision and a medical history without any psychiatric or neurological disorders. All participants provided informed written consent before the start of the experiment and received vouchers or credit points after their participation. The experiment was approved by the Ethical Committee at Maastricht University and was performed in accordance with the Declaration of Helsinki.

### Stimuli

Sixteen one-second video clips (25 frames) were used in this experiment. Each video depicted a male actor performing an emotional body movement in an angry, happy, fearful or non-emotional (e.g. coughing, pulling the nose or walking) manner. Thus, each of the four movement categories consisted of four different videos. All actors were dressed in black and their faces were blurred with a Gaussian filter to avoid triggering facial perception processes. The movements were filmed against a green background under controlled lighting conditions. The resulting clips were computer-edited using Ulead and After Effects. The videos used in this experiment were selected from a larger validated stimulus set to ensure high recognition accuracy (>80%). Detailed information regarding the recording and validation of these stimuli can be found in Kret et al. (2011b).

### Pose estimation

The state-of-the-art 2D pose estimation library OpenPose (v1.0.1, Cao et al., 2017) was used to estimate the pose of each actor in the video stimuli. By means of a convolutional neural network, OpenPose estimates the position (i.e. x- and y-coordinates) of a total of 18 *keypoints* corresponding to the main body joints (i.e. ears, eyes, nose, neck, shoulders, elbows, hands, left and right part of the hip, knees and feet). Subsequently, a skeleton is produced by association of pairs of *keypoints* using part affinity fields (see **Figure 1.A** for examples of our stimuli with the OpenPose skeleton). In the current study, the *keypoints* belonging to the eyes and ears were excluded from further analyses since the blurring of the actors’ faces often resulted in an inaccurate location estimation. The *keypoint* corresponding to the nose was kept, however, as a reference for head position. Therefore, the x- and y-coordinates were obtained for a total of 14 *keypoints,* for the 25 frames of each video in the stimulus set. Manual corrections on the estimated joint positions were performed when necessary using the coordinate system of Adobe Photoshop CS6 (v13.0, Adobe Systems Inc., San Jose, CA, USA).

**Figure 1.**
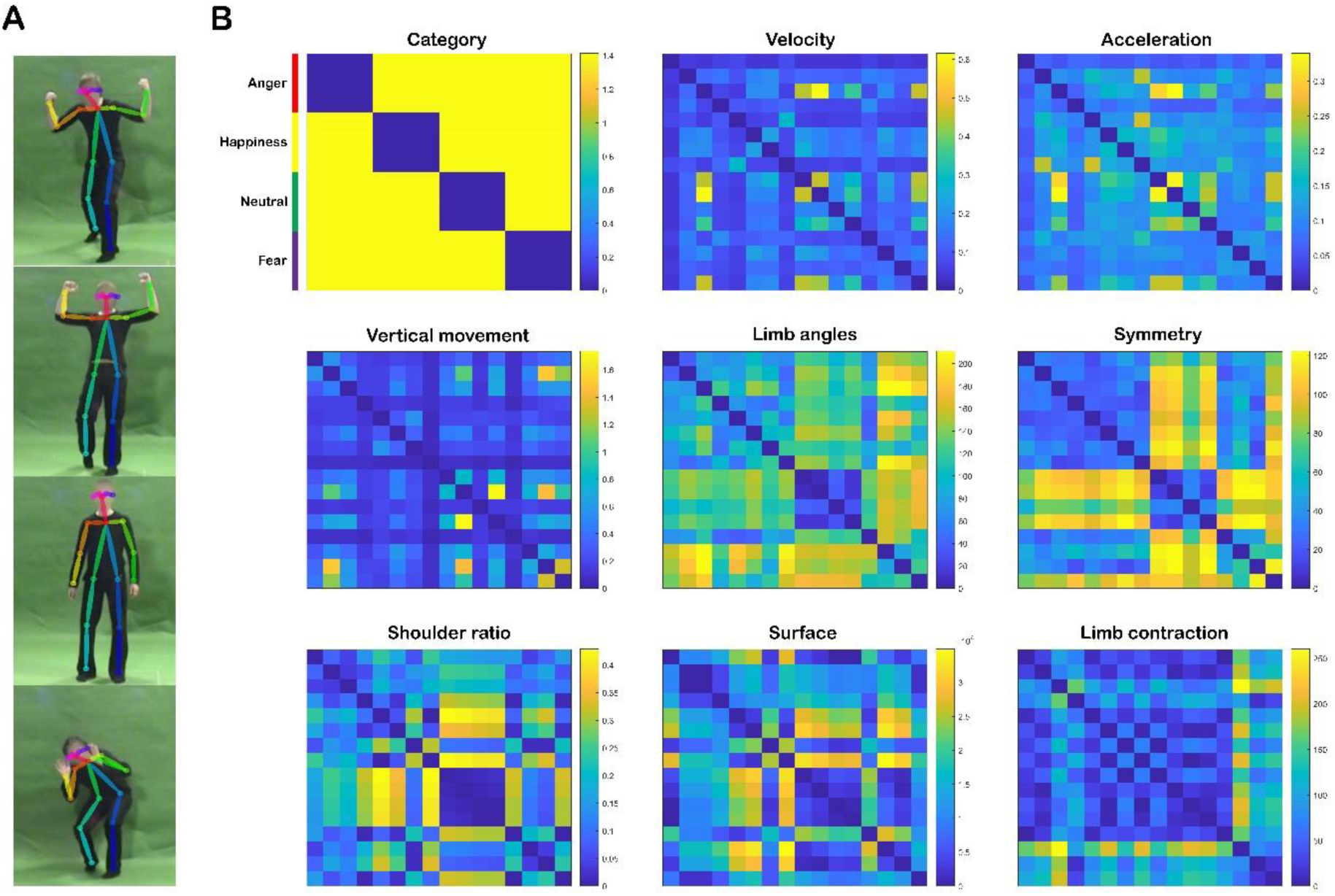
Representational dissimilarity matrices of the kinematic and postural features. **A)** Examples of frames from the different affective movement videos with the OpenPose skeleton; **B)** The RDMs represent pairwise comparisons between the 16 stimuli with regard to the kinematic (i.e. velocity, acceleration and vertical movement) and postural features (i.e. limb angles, symmetry, shoulder ratio, surface and limb contraction) averaged over time. The dissimilarity measure reflects Euclidean distance, with blue indicating high similarity and yellow high dissimilarity. Colour lines in the upper left corner indicate the organization of the RDMs with respect to the emotional category (anger: red; happiness: yellow; neutral: green; fear: purple) of the video stimuli.

### Feature definition

Several quantitative features were computed to investigate the role of body posture and kinematics in the processing of affective body movements. The features were selected due to their relevance in previous work (for a review see Kleinsmith & Bianchi-Berthouze, 2012) and included velocity, acceleration, vertical movement, symmetry, limb angles and different definitions of body contraction (i.e. shoulder ratio, surface and limb contraction). These features were computed in the same manner as in Poyo Solanas, Vaessen, & de Gelder (2020) using custom code in MATLAB (vR2017a, The MathWorks Inc., Natick, MA, USA). For an overview of the feature definition procedure, see **Table 1**. Initially, each feature was calculated for each frame; however, all values were averaged over the duration of the video clip (i.e. 25 frames) for their comparison to the imaging data.

**Table 1.** Feature definition

|  |  |  |
| --- | --- | --- |
| **Kinematic** | **Velocity** | Euclidean distance in pixel space of each *keypoint* between contiguous frames. |
|  | **Acceleration** | Difference in velocity between adjacent frames for each *keypoint*. |
|  | **Vertical movement** | Difference in y-axis pixel coordinates of each *keypoint* between adjacent frames. |
| **Postural** | **Limb angles** | Angle between two adjacent body segments, including the angles for the elbows, knees, shoulders and hips. |
|  | **Symmetry** | Euclidean distance in pixel space between each pair of joints (i.e. one on the left side, the other on the right) with respect to the axis that divides the body vertically by the nose. |
|  | **Shoulder ratio** | Amount of extension of the body joints with respect to the shoulders (measured as Euclidean distance in pixel space). |
|  | **Surface** | Surface area spanned by the total body extension in the x-axis and the extension in the y-axis (measured as Euclidean distance in pixel space). |
|  | **Limb contraction** | Average of the Euclidean distances in pixel space between the wrists and ankles to the head. |

Note: each feature was initially calculated for each frame, although the time information was later averaged.

### Experimental design, task and procedure

The (f)MRI data used in this paper was collected as part of another study. The original experiment consisted of two experimental sessions, one presenting face and voice stimuli and the other one body and voice stimuli. In each session, six experimental runs and an anatomical run were acquired. In addition, three different functional localizer runs were collected in total per subject. From this point onwards, only the stimuli, task and procedures concerning the current research goals will be described (i.e. body stimuli/runs). For a full description of the original study, see Vaessen, Van der Heijden, & de Gelder (2019).

The functional runs of the main experiment employed an event-related paradigm. Each run started with the presentation of the sixteen video clips that comprised the body movement stimulus set, followed by sixteen voice clips. Each trial started with a fixation cross, followed by the presentation of a one-second clip, with an inter-stimulus interval of 1600 – 1900ms (blank screen). In addition to the stimulus trials, each run presented four catch trials, where a video clip was presented followed by a fixation cross that changed colour to red or blue. Participants were given two seconds to press a button indicating the perceived colour. This was performed to ensure that participants were paying attention to the task while not explicitly directing their focus of attention to the explicit evaluation of the emotional expression. The total duration of each functional run was eight minutes.

The functional localizer run used one-second dynamic stimuli from the same stimulus set as the stimuli in the main experiment, but also included the corresponding dynamic faces and voices. A total of eight male actors expressing anger, happiness, fear or a non-emotional expression constituted the localizer stimuli. Some identities overlapped those used in the main experiment. The body clips displayed actors with the face blurred, dressed in black and filmed against a green background. For the facial videos, actors wore a green shirt similar to the background colour. All faces were recorded from the frontal view and did not include any information below the neck. The auditory stimuli consisted of non-verbal vocalizations. These stimuli were combined giving a total of six audio-visual and unimodal stimuli categories: (1) face alone, (2) body alone, (3) voice alone 1, (4) voice alone 2, (5) face and voice, and (6) body and voice. Two sets of isolated non-verbal vocalizations (i.e. voice alone 1 & 2) were presented to have similar statistical power when comparing conditions. These two conditions differed in the stimuli used. Each category was presented ten times using a block design under passive viewing conditions. In each block, eight stimuli of the same category were presented in random order, with two different stimuli per emotional expression. The total duration of each block was eight seconds. Each block was followed by an eight-second fixation period. The total duration of the localizer run was approx. 16 min.

Visual stimuli were displayed using Presentation software (v19.0, Neurobehavioral Systems, Inc., Berkeley, USA) and back-projected with a LCD projector onto a screen (screen resolution = 1920 x 1200; screen height = 24.5 cm; screen width = 40 cm; screen diagonal = 47 cm) situated at the posterior end of the scanner bore. Participants viewed the stimuli through a mirror attached to the head coil (screen–mirror distance = 60 cm; mirror–eye distance = 15 cm approximately; total screen–eye distance = 75 cm approximately). The video clips had a resolution of 1080 x 864 pixels (visual angles = 8.5 x 6.7 degrees). Auditory stimuli were presented through MR-compatible, sound-attenuating earplugs.

### (f)MRI data acquisition

Data were acquired with a 3 Tesla whole-body scanner (Siemens, Erlangen, Germany) located at the Maastricht Brain Imaging Centre (MBIC) of Maastricht University, the Netherlands. Functional images of the whole brain were obtained using T2*-weighted 2D echo-planar image (EPI) sequences [number of slices per volume = 50, 2 mm in-plane isotropic resolution, repetition time (TR) = 3000 ms, echo time (TE) = 30 ms, flip angle (FA) = 90°, field of view (FoV) = 800 x 800 mm^2^, matrix size = 100 x 100, multi-band acceleration factor = 2, number of volumes per run = 160, total scan time per run = 8 min]. The functional localizer scan also used a T2*-weighted 2D EPI sequence [number of slices per volume = 64, 2 mm in-plane isotropic resolution, TR = 2000 ms, TE = 31 ms, FA = 77, FoV = 800 x 800 mm^2^, matrix size = 100 x 100, multi-band acceleration factor = 2, number of volumes per run = 491, total scan time per run = 16 min approx.]. Three-dimensional (3D) T1-weighted (MPRAGE GRAPPA2) imaging sequences were used to acquire high-resolution structural images for each participant (1-mm isotropic resolution, TR = 2250 ms, TE = 2.21 ms, FA = 9°, matrix size = 256 x 256, total scan time = 7 min approx.).

### Data pre-processing and analysis

#### (f)MRI data pre-processing

BrainVoyager (v21.2 Brain Innovation B.V., Maastricht, the Netherlands) as well as custom code in MATLAB (vR2017a, The MathWorks Inc., Natick, MA, USA) were used for the pre-processing and analysis of the acquired (f)MRI data. The pre-processing of the functional data comprised several steps. Trilinear/sinc estimation and interpolation were applied to correct for participant’s 3D head motion with respect to the first volume of each functional run. Sinc interpolation was used to correct for time differences in slice acquisition order within one volume. High-pass temporal filtering was employed to exclude low frequency drifts in the data lower than two cycles per run. The functional data of the main experiment was not spatially smoothed to preserve spatial specific information for the multivariate analyses. Spatial smoothing was applied, however, to the functional localizer data with a Gaussian kernel of a full-width half-maximum of 3 mm. The anatomical data was corrected for B1-field inhomogeneities. After these steps, the native functional and anatomical data were co-registered and template-based normalized to Talairach space (Talairach & Tournoux, 1988).

#### HRF estimation

In order to account for possible differences in the temporal delay of the blood-oxygen-level dependent (BOLD) response across the brain, an optimized hemodynamic response function (HRF) model was used (De Angelis et al., 2018). For this purpose, the unsmoothed pre-processed time courses of each individual subject were divided, after regressing out the motion parameters, into separate trials according to each stimulus presentation time. Each event was defined as a six TR window (i.e. comprising the TR before stimulus onset and the four TRs after), which was percentage-signal-change normalized. Subsequently, the normalized BOLD response of all trials was fitted to an HRF. An optimal delay for the HRF was chosen for each voxel provided by the time-to-peak with the best fit to the trial average response, estimated by varying this parameter between 4.0s and 6.0s in 0.5s steps. As a result of this method, a beta value for each trial and voxel was obtained.

#### Region of Interest definition

The functional localizer data were used to identify several regions of interest (ROI) for body perception. For this purpose, a fixed-effects whole-brain general linear model was fitted to the 3-mm-smoothed localizer data of each participant. The generated regression model consisted of the %-signal-transformed predictors of each stimulus category (i.e. body alone, face alone, voice alone 1, voice alone 2, body-voice, face-voice) convolved with a two-gamma HRF and the z-transformed motion parameters as predictors of no interest.

The considered ROIs include occipito-temporal areas that have previously shown a certain level of body specificity (three ROIs: FBA, EBA, pSTS) (Downing et al., 2001; Kontaris, Wiggett, & Downing, 2009; Peelen & Downing, 2005; Schwarzlose et al., 2005; Vangeneugden, Peelen, Tadin, & Battelli, 2014), parietal and temporal areas thought to be implicated in attention and action observation (six ROIs: V7/3a, superior parieto-occipital cortex (SPOC), superior marginal gyrus (SMG), posterior intraparietal sulcus (pIPS), medial intraparietal sulcus (mIPS), anterior intraparietal sulcus (aIPS)) (Caspers et al., 2010; Corbetta, Patel, & Shulman, 2008; Culham & Valyear, 2006; Grafton & Hamilton, 2007), and frontal areas involved in action observation and other higher cognitive functions (six ROIs: ventral and dorsal premotor cortex (PMv, PMd), SMA, pre-SMA, inferior frontal and frontal regions) (Caspers et al., 2010; Grafton & Hamilton, 2007). Body movement videos were contrasted to baseline at uncorrected p = 0.005 to define the ROIs with the exception of FBA and EBA. For the latter, body movements were contrasted against facial movement clips at uncorrected p = 0.005. The ROIs were defined bilaterally whenever possible and subsequently merged into a single ROI for each participant individually. For a detailed explanation of the definition and location of the ROIs, see Supplementary Materials.

### Representational similarity analysis

Representational similarity analyses (RSA) (Kriegeskorte et al., 2008; Nili et al., 2014) were performed in MATLAB (vR2017a, The MathWorks Inc., Natick, MA, USA) to investigate the relations among the computed features and brain activity. This type of analysis is based on the determination of representational dissimilarities between pairs of stimuli values. The representation is characterized by symmetrical matrices called representational dissimilarity matrices (RDM). In these matrices, off-diagonal values reflect the dissimilarity between the values of two different stimuli while diagonal entries represent comparisons between identical stimuli and are zero by definition.

#### Computed-feature RDMs

RDMs were constructed based on the dissimilarity between all stimulus pairs, in Euclidean distance, with respect to the computed feature values (i.e. first-level feature RSA). Dummy variables were used to compute the emotional categories RDM, where the same emotion was defined as having zero dissimilarity with itself while two different emotions had a dissimilarity of √2. This analysis resulted in 16 x 16 distance matrices, one for each feature. To examine possible correlations among features, Spearman’s rank correlations were performed.

#### Neural RDMs

In order to create neural RDMs for both the whole-brain searchlight and the ROI analyses, the β-values of each of the 16 body stimuli presented in the main experiment were used. These β-values were obtained after the application of an optimized HRF model to the data (see HRF estimation section above). For each ROI, a first-level RSA analysis was performed consisting of comparisons between the β-values for each pair of stimuli using Pearson’s correlation r (r = 1: perfect correlation; r = -1: perfect anti-correlation). Each possible stimulus-pair distance (d) was defined by d = 1 - r, where d ranges from 0 to 2. This generated a 16 x 16 distance matrix for each ROI and participant. In a second-level RSA analysis, every ROI dissimilarity matrix was correlated to each feature RDM using Spearman’s rank correlation. The resulting correlation values were then z-transformed for each participant and a group-level one-sample t-test against 0 (2-tailed; Benjamini-Hochberg false discovery rate (BHFDR)-adjusted p-values) was computed for each feature.

A whole-brain searchlight (radius = 5 voxels) analysis was performed using custom in-house MATLAB scripts. This approach allowed us to identify areas involved in the perception of emotional body movements that were not covered by the defined ROIs. Similar to the ROI analyses, searchlight RDMs were correlated with each feature RDM. The resulting maps were z-transformed for each participant. Subsequently, a group-level one-sample t-test against 0 (2-tailed, cluster size corrected with Monte-Carlo simulation, alpha level = 0.05, initial p = 0.005, numbers of iterations = 5000) was computed for each feature. A cluster showing a positive correlation means that a high dissimilarity between a stimulus pair in the feature RDM also has a high dissimilarity in the neural representation. A negative correlation means that a low dissimilarity between a stimulus pair would have a higher dissimilarity in the neural representation. The anatomical labelling of the resulting clusters was performed according to the atlas of Duvernoy (Duvernoy, 1999) for a more reliable localization. Univariate results are not reported in this paper, but see Vaessen et al. (2019) or **Figure SR3** in Supplementary Results.

### Comparison between postural and kinematic feature processing

Paired-sample t-tests were conducted to investigate whether the pre-selected ROIs processed kinematic and postural information differently. For each subject, we averaged the correlation values obtained for each feature-ROI comparison, separately for the postural features (i.e. shoulder ratio, surface, limb distances, symmetry and limb angles), and kinematic ones (i.e. velocity, acceleration and vertical movement). Subsequently, a paired-sample t-test was conducted per ROI comparing kinematic and postural values.

### Comparison between dorsal and ventral processing of body features

We also investigated whether there was a difference in individual feature processing in dorsal and ventral clusters. For each subject, the correlation values obtained for each feature-ROI comparison were averaged within the dorsal (i.e. aIPS, mIPS, pIPS, V7/3a, SPOC, SMG and pSTS) and ventral (i.e. EBA and FBA) ROIs. Next, a paired-sample t-test was conducted per feature.

### Classification of emotional categories from fMRI data

A Gaussian Naïve Bayes (GNB) classifier (Ontivero-Ortega, Lage-Castellanos, Valente, Goebel, & Valdes-Sosa, 2017) was used to classify the emotional category from the fMRI data. For each participant, the classifier was trained and tested with a leave-one-run-out cross-validation procedure using the searchlight data (see the *Neural RDMs* section above) as input. Subsequently, a group-level one-sample t-test against chance level (i.e. 25%) was performed with the resulting decoding accuracies.

## Results

### Kinematic and postural features

Representational similarity analyses were carried out to examine whether the defined features reflected the affective categorical structure of the body stimuli. This same analysis was performed in a behavioural study with a larger stimulus set that included the current 16 video clips (Poyo Solanas et al., 2020). A first inspection of the RDMs revealed that kinematic body features (i.e. velocity, acceleration and vertical movement) do not clearly represent the categorical structure of emotion (see **Figure 1.B**). Generally, these features showed relatively high similarity between categories. Only velocity and acceleration showed a distinctive level of within-category similarity for the neutral condition. High within-category similarity was also found for neutral in the majority of postural features. In the case of symmetry, this finding was also accompanied by a high level of dissimilarity between neutral and the rest of the affective movement categories and a within- and between-category similarity for anger and happiness. The neutral and fearful conditions were different from the happy and angry categories for the features limb angles, shoulder ratio and surface. Limb contraction was relevant for differentiating between fear and the rest of the categories.

Pairwise comparisons between kinematic and postural matrices were performed to examine their interrelation and their relation to emotional categories (**Figure 2**; see **Table SR1** in Supplementary Results for correlation and p-values). While kinematic features did not show significant correlations to emotion categories, postural features revealed week to moderate significant correlations. Specifically, limb angles (r(118) = 0.408, pBonf-corr. < 0.001), symmetry (r(118) = 0.349, pBonf-corr. = 0.001) and shoulder ratio (r(118) = 0.345, pBonf-corr. = 0.001) showed the strongest correlations to emotion. The relationship between postural and kinematic features was weak and often negative. Kinematic features correlated strongly among themselves, and similar findings were observed among postural features.

**Figure 2.**
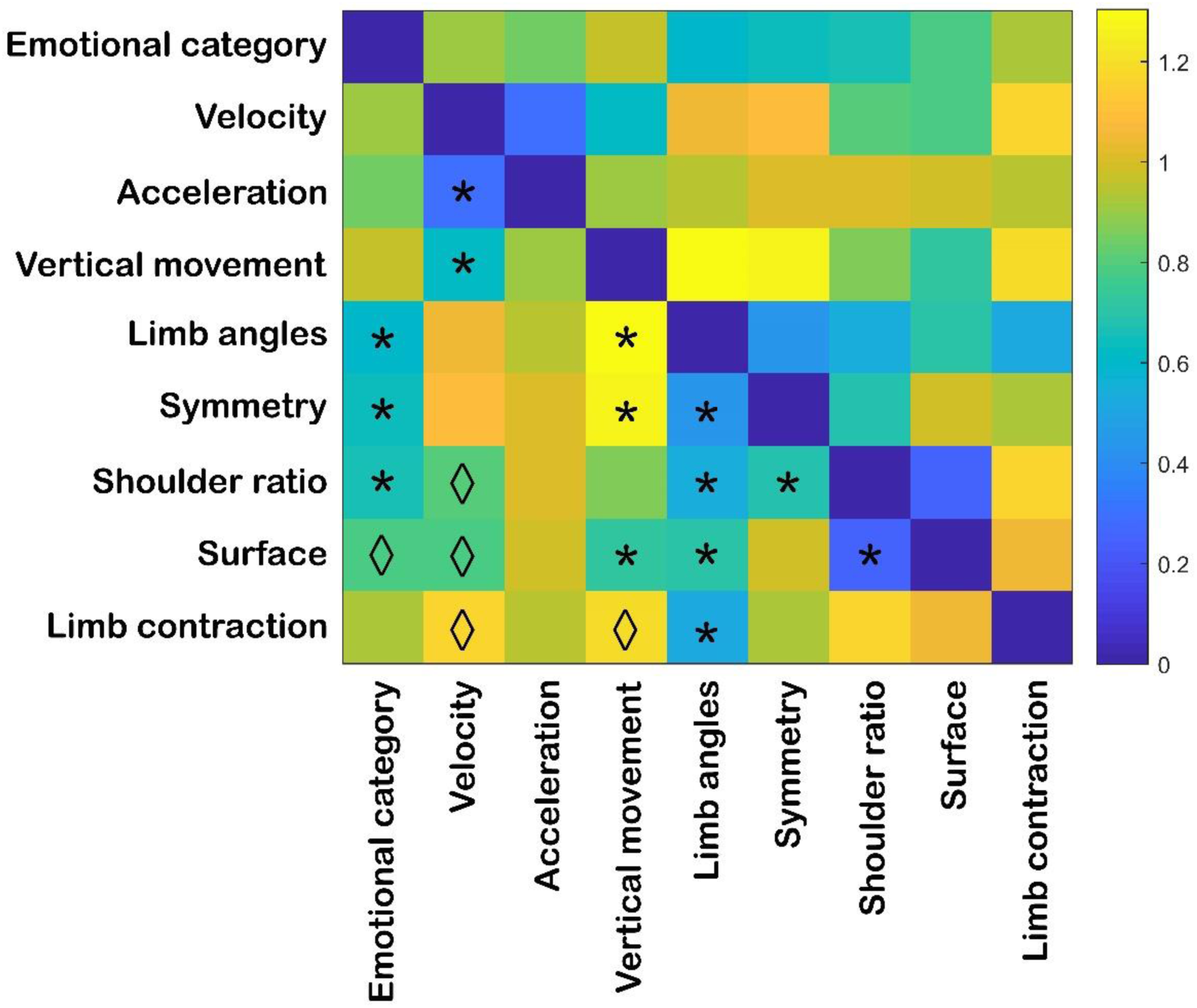
Correlation between representational dissimilarity matrices of kinematic and postural features. The RDM represents the level of (dis)similarity between each of the kinematic (i.e. velocity, acceleration and vertical movement) and postural (i.e. limb angles, symmetry, shoulder ratio, surface and limb contraction) matrices (see Figure 1). Distances are indicated in 1-Spearman’s correlation values, with blue indicating high similarity and yellow high dissimilarity. Asterisks and rhombi below the diagonal indicate significant correlations after Bonferroni correction and correlations that presented significant uncorrected p-values, respectively (αbonf = 0.05/9, with nine comparisons per feature; see Table SR1 in Supplementary Results for correlation and p-values).

### Kinematic and postural feature representation in body-selective areas

The second aim of the study was to determine whether (dis)similarities in body posture and/or body-movement kinematics between different emotional categories could explain the neural response at the whole-brain level and in body-selective regions of interest. Several areas were defined known for their involvement in the processing of body expressions and their neural RDMs were computed (see Methods and Supplementary Materials for more information regarding ROI definition and location. For an inspection of the ROI matrices see **Figure SR1**). Subsequently, each neural matrix was correlated to each feature RDM and a group-averaged correlation value was obtained per feature-ROI combination (see **Figure 3**).

**Figure 3.**
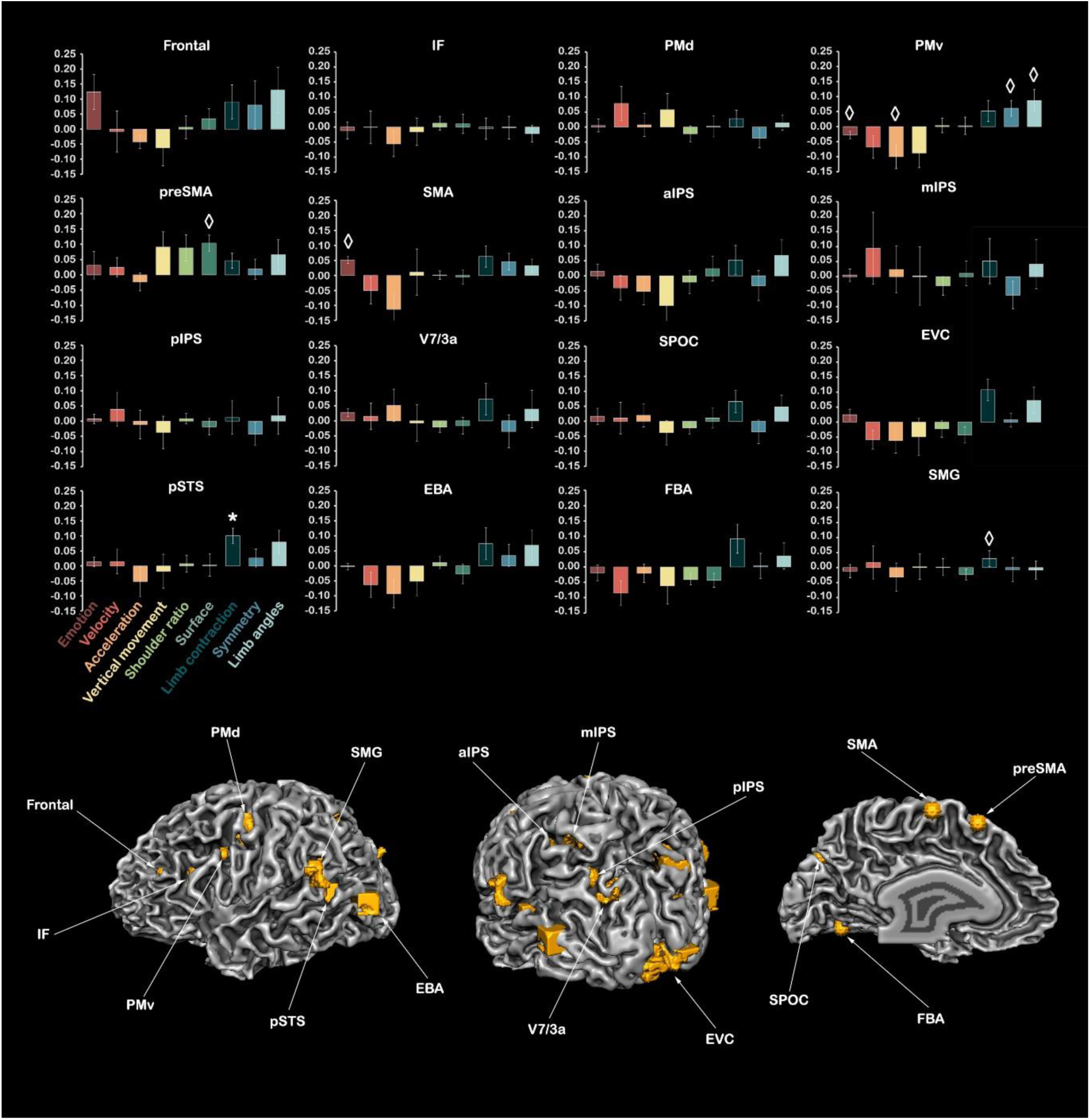
Average Spearman’s rank correlation across participants between the kinematic/postural feature RDMs and each ROI matrix. Kinematic features include velocity, acceleration and vertical movement. Postural features comprise shoulder ratio, surface, limb contraction, symmetry and limb angles. Positive r values indicate that a high dissimilarity between a stimulus pair in the feature RDM also has a high dissimilarity in the neural representation. A negative correlation means that a low dissimilarity between two stimuli at the feature level would have a higher dissimilarity in the neural representation. Asterisks and rhombi indicate significant correlations after BHFDR correction and correlations that presented significant uncorrected p-values, respectively (one sample t-test against 0, two-tailed). The error bars denote the standard error of the mean (SEM). Order or relationships across ROIs are not assumed here. Abbreviations: EBA: extrastriate body area; EVC: early visual cortex; FBA: fusiform body area; IF: inferior frontal gyrus; IPS: intraparietal sulcus, p: posterior, m: middle, a: anterior; PMd: dorsal premotor cortex; PMv: ventral premotor cortex; pre-SMA: pre-supplementary motor area; pSTS: posterior superior temporal sulcus; SMA: supplementary motor area; SMG: supramarginal gyrus; SPOC: superior parietal occipital cortex.

The results of this analysis showed that PMv (negatively, r(10) = -0.03, p_uncorrected_ = 0.047, p_Bonferroni corrected_ = 0.349) and SMA (positively, (r(6) = 0.05, p_uncorr._ = 0.004, p_Bonf-corr._ = 0.066) correlated significantly to emotion categories, but only before correction for multiple comparisons. Although non-significant, the relatively highest positive correlation to emotion categories was found in the frontal cluster (r(3) = 0.12, p_Bonf-corr._ = 0.500). The features representing kinematic features of body movement mainly displayed negative correlations to the majority of the defined ROIs. However, only the negative correlation between PMv and acceleration was significant before correction for multiple comparisons (r(10) = -0.10, p_uncorr._ = 0.031, p_Bonf-corr._ = 0.397). Though not significant, the relatively highest positive correlations to the kinematic features were found in pSTS, pIPS and EBA, although the latter one only for vertical movement. With regard to postural features, a significant positive correlation was found between pSTS and limb contraction (r(11) = 0.10, pBonf-corr. = 0.038). At uncorrected p-value, also between PMv and symmetry (r(10) = 0.06, p_uncorr._ = 0.043, p_Bonf-corr._ = 0.689), PMv and limb angles (r(10) = 0.09, p_uncorr._ = 0.044, p_Bonf-corr._ = 0.441), pre-SMA and surface (r(4) = 0.10, p_uncorr._ = 0.017, p_Bonf-corr_ = 0.276) and EVCF and limb contraction (r(12) = 0.11, p_uncorr._ = 0.011, p_Bonf-corr._ = 0.086).

Next, pair-sample t-tests were performed to examine whether the pre-selected ROIs processed kinematic and postural information differently. Only PMv showed a significant difference in the processing of kinematic (M = -0.09, SD = 0.08) and postural (M = 0.04, SD = 0.05) features at uncorrected p-value; t(10) = -3.8, p_uncorr._ = 0.004, p_Bonf-corr._ = 0.056 (see **Table SR2** in Supplementary Results for an overview of all paired-sampled t-tests). In addition, we investigated whether there was a difference in the processing of each individual feature with regard to dorsal and ventral ROIs. Only velocity was processed significantly different in dorsal (M = 0.03, SD = 0.15) and ventral ROIs (M = -0.06, SD = 0.15) at uncorrected p-value; t(12) = -2.23, p_uncorr._ = 0.046, p_Bonf-corr._ = 0.411 (see **Table SR3** in Supplementary Results for an overview of all paired-sampled t-tests).

To investigate whether the defined ROIs share or present unique affective body-movement representations, the group-averaged neural matrix of each ROI was correlated to that of the other ROIs. Overall, there was a relatively high similarity between and within parietal and temporo-occipital regions (see **Figure 4**; see **Table SR4** in Supplementary Results for correlation and p-values). Specifically, SMG and EBA (r(1538) = 0.64, pBonf-corr < 0.001), SMG and pSTS (r(1538) = 0.58, pBonf-corr < 0.001), pIPS and EBA (r(1538) = 0.57, pBonf-corr < 0.001), pSTS and EBA (r(1538) = 0.57, pBonf-corr < 0.001) and pIPS and SMG (r(1538) = 0.47, p_Bonf-corr_ < 0.001) showed the strongest significant positive correlations. The inferior frontal cluster and PMv also showed significant positive correlations with temporo-parietal areas. Although no significant negative correlations were found after Bonferroni correction, frontal and premotor regions presented some degree of dissimilarity, as well as frontal and ventral areas, and motor and parietal regions.

**Figure 4.**
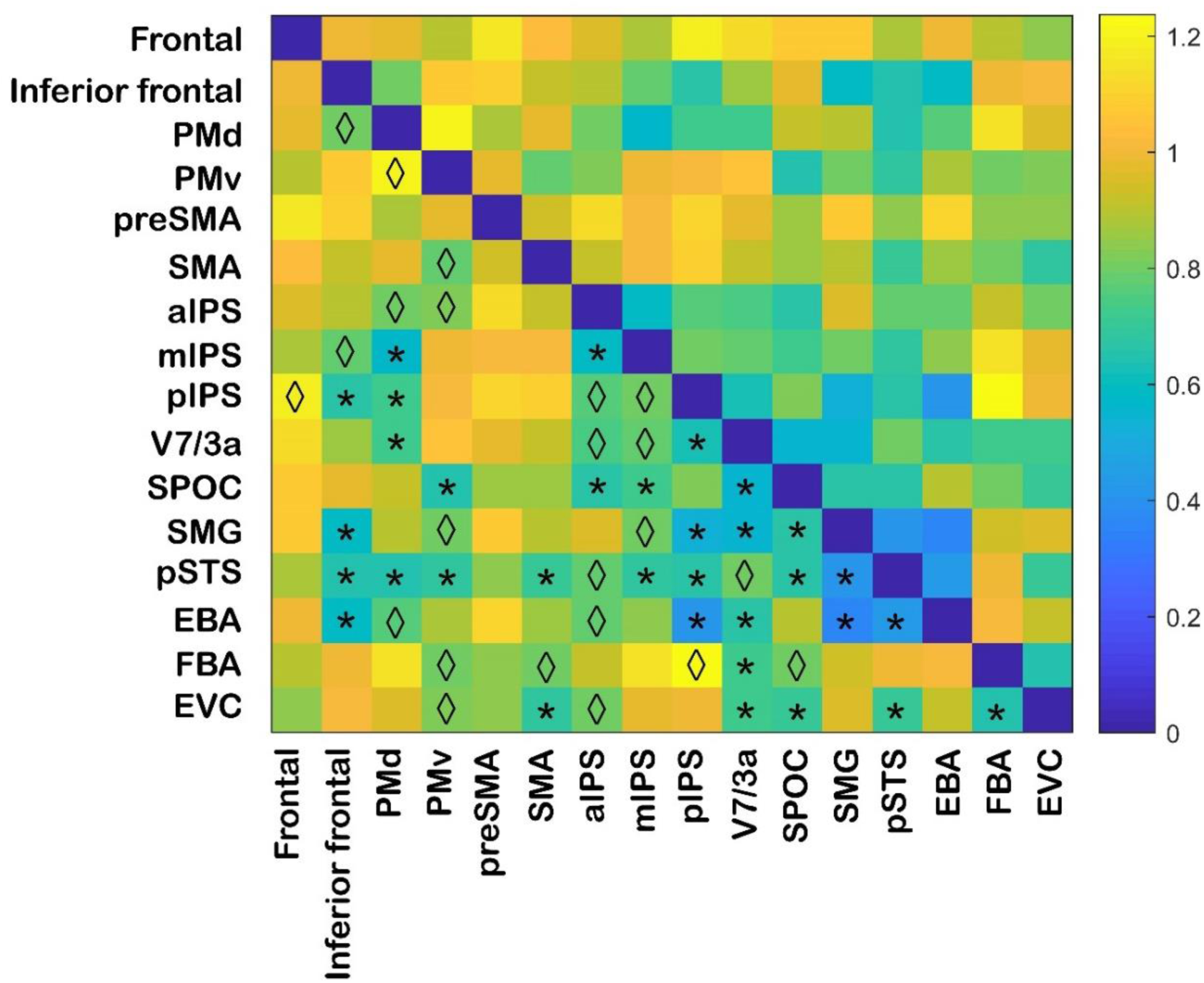
Correlation between ROI representational dissimilarity matrices. For each ROI, a group-averaged RDM was obtained. Subsequently, pairwise comparisons between the resulting group ROI matrices were performed. The dissimilarity measure reflects 1-Spearman’s rank correlation values, with blue indicating strong similarity and yellow strong dissimilarity. Rhombi and asterisks below the diagonal indicate significant correlations before and after Bonferroni correction for multiple comparisons, respectively (αbonf = 0.05/16, with 16 comparisons per ROI, see Supplementary Results Table SR4). Abbreviations: EBA: extrastriate body area; EVC: early visual cortex; FBA: fusiform body area; IPS: intraparietal sulcus, p: posterior, m: middle, a: anterior; PMd: dorsal premotor cortex; PMv: ventral premotor cortex; pre-SMA: pre-supplementary motor area; pSTS: posterior superior temporal sulcus; SMA: supplementary motor area; SMG: supramarginal gyrus; SPOC: superior parietal occipital cortex.

### Whole-brain kinematic and postural feature representation with multivariate approaches

In addition to the body-selective regions, we also investigated whether (dis)similarities in body posture and/or body-movement kinematics between different emotional categories could explain the neural response at the whole-brain level. The computed feature RDMs were compared to the multi-voxel dissimilarity fMRI patterns by means of searchlight RSA. The clusters resulting from this analysis are shown in **Figure SR2** and **Table SR5** in Supplementary Results. The matrix for velocity was positively correlated to inferior frontal sulcus and precentral gyrus. Negative main effects for acceleration were found in middle temporal, superior frontal and postcentral sulci while no positive main effects were observed for this feature. Vertical movement correlated positively with cingulate gyrus, whereas negatively to the frontomarginal and middle temporal gyri.

With respect to postural features, limb angles showed a positive main effect in anterior insula and pSTS. Several areas negatively correlated to symmetry in the inferior and middle occipital gyri, precuneus, isthmus, anterior calcarine, intraparietal and cingulate sulci. Shoulder ratio negatively correlated to anterior insula, frontal operculum, putamen, ACC, middle frontal gyrus, cingular insular sulcus, claustrum, internal capsule and parahippocampal gyrus. Surface showed main negative effects in posterior orbital gyrus, thalamus, anterior perforated substance, ACC, inferior and superior frontal sulci, putamen and internal capsule. Only positive correlations to limb contraction were found in intraparietal sulcus, anterior insula, caudate nucleus, amygdala, superior frontal sulcus and gyrus, precuneus, posterior orbital gyrus, ACC, superior temporal gyrus, inferior precentral sulcus and SMG.

### Whole-brain representation of emotion with multivariate approaches

We searched for representations of emotion at the whole-brain level by running a searchlight RSA with the emotion category RDM. No area showed a main effect of emotion after correction for multiple comparisons. Before correction, however, several regions correlated positively to emotion, including inferior temporal cortex, anterior insula, lingual sulcus, superior frontal gyrus and lateral occipital sulcus (see **Table SR6** in Supplementary Results for an overview of the clusters showing a main effect of emotion at uncorrected p-value). Interestingly, negative main effects of emotion were found in amygdala, inferior frontal and middle temporal gyri. In addition to the RSA analysis described above, we also ran an analysis where a GNB classifier was trained to decode the emotion of the stimuli from the unsmoothed multi-voxel brain patterns. The clusters resulting from the GBN classification of emotion were different to those from the RSA (see **Figure 6** and **Table SR7** in Supplementary Results). The areas showing above chance accuracy included ACC, pSTS, middle temporal sulcus, inferior frontal, angular, supramarginal, fusiform and inferior temporal gyri.

## Discussion

The current study aimed at investigating the relation between features defining body posture and kinematics and their brain representation. For this purpose, quantitative features of body form and motion were computed from videos of affective body movements. Representational similarity analyses were performed to investigate whether these features reflect the emotional categorical structure. By means of multi-voxel pattern analysis techniques, we also examined whether the (dis)similarity of body posture and kinematics between different emotional classes could explain the neural response in and beyond body-selective regions.

The results showed that form rather than motion-related features represent better the conveyed affect. Moreover, body movements differentially activated brain regions based on their postural and kinematic characteristics, indicating that these aspects might be encoded in these regions. Among these features, the degree of limb contraction seems to be particularly relevant for distinguishing fear from other affective movements. This feature was represented in several regions spanning affective, action observation and motor preparation networks.

### The role of kinematic and postural features in emotion recognition

In the representational similarity results, kinematic features revealed high similarity across emotions and weak correlations to the emotional categories while postural cues presented clearer distinctions (see **Figure 1**). This suggests that postural rather than kinematic body features reflect the affective categorical structure of the body movements. These results are consistent with the findings from a larger study (Poyo Solanas et al., 2020). The importance of postural information for affective recognition has been supported by previous literature (for a review see Kleinsmith & Bianchi-Berthouze, 2012). For example, Atkinson and colleagues (2007) reported that while motion cues can be sufficient for emotional recognition, the disruption of form information disproportionately compromises the recognition performance, especially in the case of fear.

The majority of postural features clearly differentiated emotional from non-emotional body movements, such as limb angles, shoulder ratio and surface. However, this distinction was most clear in the symmetry RDM, indicating that emotional movements are less symmetrical than non-emotional ones (Poyo Solanas et al., 2020). Limb angles, shoulder ratio and surface also showed dissimilarities in the representation of happy and anger from fear and neutral expressions. This could be due to the degree of openness of the body, higher for angry and happy expressions and smaller for neutral and fearful ones (for a review see Kleinsmith & Bianchi-Berthouze, 2012). In line with previous work, the degree of limb contraction seems to be relevant for distinguishing fear from other affective movements (Poyo Solanas et al., 2020; Roether et al., 2009).

### The representation of kinematic and postural features in body-selective areas

According to the two-stream model of visual processing (Giese & Poggio, 2003; Milner & Goodale, 2006, 2008; Vaina et al., 1990), form and movement information are processed in two separate pathways in the brain. In the current paper, however, no clear dorsal vs. ventral stream segregation was found with regard to the processing of kinematic and postural features in pre-selected body-selective regions. The only kinematic feature that was represented differently in ventral and dorsal areas was velocity (see Results and **Table SR3** in Supplementary Results). Particularly, dorsal regions processed body movements with comparable velocity characteristics in a similar manner and regardless of the conveyed affect, whereas ventral areas represented body movements differently despite presenting similar velocity values. These findings are indeed in agreement with the hypothesis that the dorsal stream is specialized in processing motion signals irrespective of visual forms while ventral areas use structural information rather than kinematic cues to differentiate between stimuli (Giese & Poggio, 2003; Milner & Goodale, 2006, 2008; Vaina et al., 1990).

The processing of kinematic and postural features, respectively, was significantly different in one region: PMv (see Results and **Table SR2** in Supplementary Results). This area processed body movements with comparable symmetry and limb-angle values in a similar manner, irrespective of their kinematic characteristics in terms of acceleration (see **Figure 3** and **Table SR2** in Supplementary Results). This finding is in disagreement with previous TMS studies suggesting the involvement of this area in the processing of action signals regardless of visual forms (Candidi et al., 2011; Urgesi, Candidi, Ionta, & Aglioti, 2007). The pattern of (dis)similarities observed in the RDMs of these features suggests that PMv processed neutral and fearful body movements differently from the rest of the emotion categories (see **Figure 1**). Although its role in emotional discrimination is still controversial (Candidi et al., 2011), and indeed the affective structure of the stimulus set was not well represented by this area’s activity pattern, PMv seemed to distinguish between threatening and neutral expressions, in agreement with previous neuroimaging (Calbi, Angelini, Gallese, & Umiltà, 2017; Kret et al., 2011b; Pichon, de Gelder, & Grèzes, 2008) and TMS studies (Balconi & Bortolotti, 2013; Engelen, Zhan, Sack, & de Gelder, 2018). This is an interesting finding since PMv has been involved in action preparation and execution (Hoshi & Tanji, 2007; Picard & Strick, 2001) as well as in space perception and action understanding (Rizzolatti, Fadiga, Gallese, & Fogassi, 1996; Rizzolatti, Fogassi, & Gallese, 2002; Urgesi, Calvo-Merino, Haggard, & Aglioti, 2007; Urgesi, Candidi, et al., 2007).

The posterior superior temporal sulcus, an area known to be involved in the processing of biological motion (Allison, Puce, & McCarthy, 2000; Grossman et al., 2000; Grossman et al., 2010), did not seem to encode kinematic body features. However, this area processed body movements with similar limb contraction characteristics in a similar manner (see **Figure 3**).

From the stimuli representation in the limb contraction RDM, this finding suggests that pSTS may process fearful body movements in a dissimilar manner to the rest of the affective categories (see **Figure 1**). This is in line with previous studies showing the involvement of this area in the recognition of emotions (Basil, Westwater, Wiener, & Thompson, 2017; Wegrzyn et al., 2015; Zhang et al., 2016), especially when the expression conveyed depicts fear (Candidi et al., 2011; Grèzes et al., 2007).

The pre-supplementary motor area showed a positive correlation to the representation of surface (see **Figure 3**). This result indicates that pre-SMA may be relevant for the discrimination of neutral and fearful expressions from happy and angry ones (see **Figure 1**). This area may use this structural body feature to understand the intention behind the observed movement (Lau, Rogers, Haggard, & Passingham, 2004) and prepare for an appropriate motor response (Isoda, 2005; Luppino, Matelli, Camarda, Gallese, & Rizzolatti, 1991). Finally, EBA and FBA did not represent the kinematic aspects of the affective movements but showed greater tuning to postural cues, although not consistently or reaching significance (see **Figure 3**). This is in line with previous literature involving these areas in the processing of bodies and body parts (Peelen & Downing, 2007). However, the stimuli representation in EBA was very dissimilar to that of FBA (see **Figure 4** and **Table SR4** in Supplementary Results), which may reflect their different roles in body processing. It has been suggested that EBA represents separate body parts while FBA is more involved in the processing of whole bodies (Hodzic, Kaas, Muckli, Stirn, & Singer, 2009; Taylor, Wiggett, & Downing, 2007; Urgesi, Calvo-Merino, et al., 2007). Interestingly, the results also showed that SMG, pSTS, pIPS and the inferior frontal cortex may represent body movements in a similar manner to EBA while dissimilarly to FBA (see **Figure 4** and **Table SR4** in Supplementary Results).

### Whole-brain kinematic and postural feature representation with multivariate approaches

Both cortical and subcortical areas were recruited for the processing of kinematic and postural features (see **Figure 5**). As in the results following the region of interest analyses, no clear dorsal vs. ventral stream dissociation was observed for the processing of these features. There was no overlap among brain regions correlating to kinematic descriptors and the stimuli representation in these areas was often dissimilar to that of the kinematic feature RDMs, which presented an overall high similarity between and within emotional states (see **Figure 1**). One of these regions, pSTS, correlated negatively to the displacement of the body joints in the vertical axes (see **Figure 5** and **Table SR5** in Supplementary Results). This suggests that this area may be able to discriminate between different movements despite presenting similar vertical displacement of the body joints, and regardless of the emotion conveyed. The previously discussed results on pre-selected ROIs showed that pSTS might represent fearful body movements differently to the rest of the affective categories, and that it may use the information conveyed by the contraction of the limbs to perform this discrimination (see **Figure 1** and **Figure 3**). Further research will be needed to clarify whether pSTS is modulated by emotion and to understand how and which low-level visual features this area uses for the discrimination between affective body movements.

**Figure 5.**
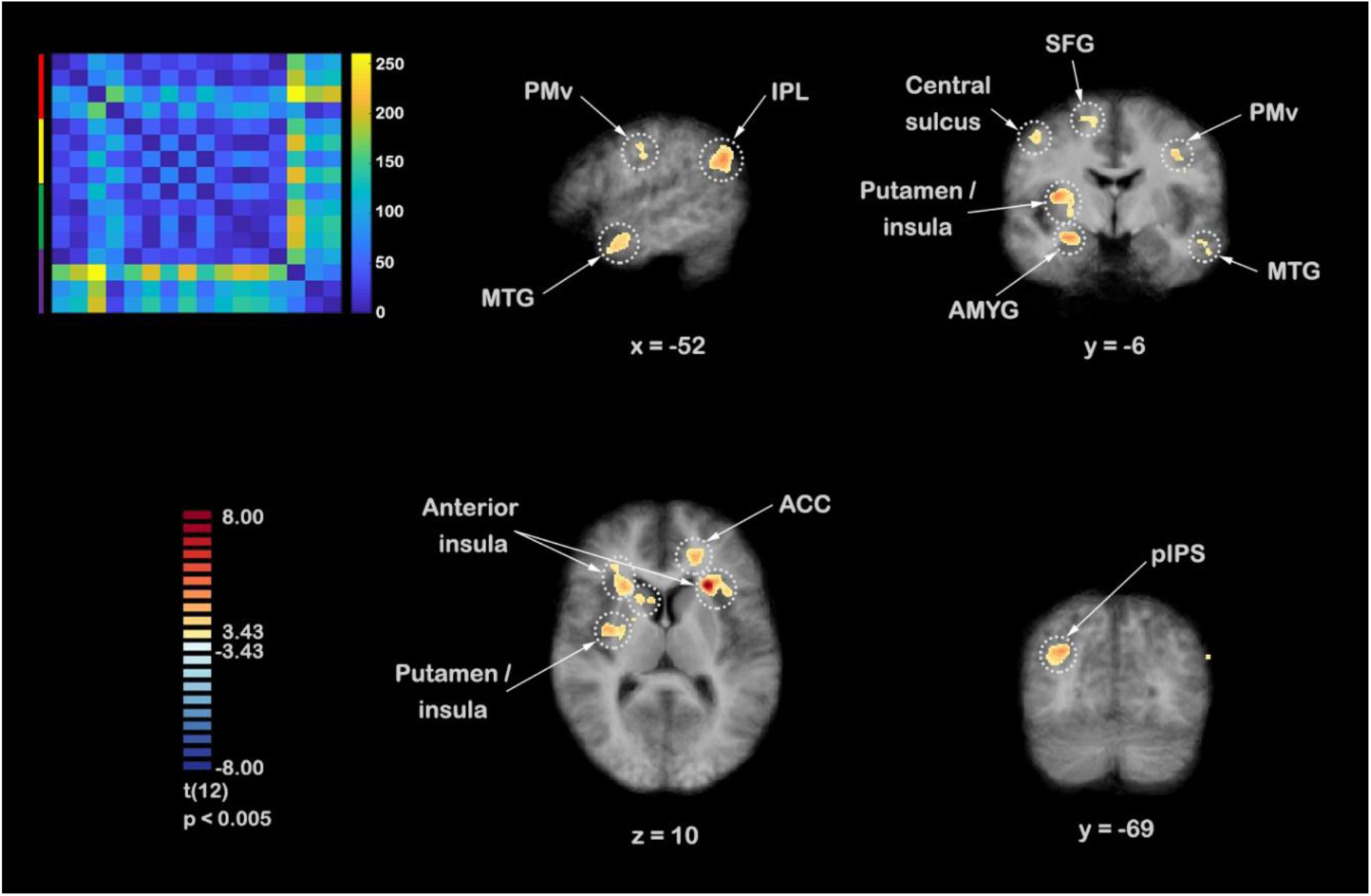
Clusters resulting from the searchlight RSA of the postural feature of limb contraction. The multi-voxel fMRI dissimilarity matrices were correlated to the limb contraction RDM (upper left corner). The resulting maps were z-transformed for each participant. Subsequently, a group-level one-sample t-test against 0 was performed (2-tailed, cluster size corrected with Monte-Carlo simulation, alpha level = 0.05, initial p = 0.005, numbers of iterations = 5000). See Table SR5 in Supplementary Results for more details on location and statistical values of the clusters. Abbreviations: ACC: anterior cingulate cortex; AMYG: amygdala; IPL: inferior parietal lobule; MTG: middle temporal gyrus; pIPS: posterior intraparietal sulcus; PMv: ventral premotor cortex; SFG: superior frontal gyrus.

**Figure 6.**
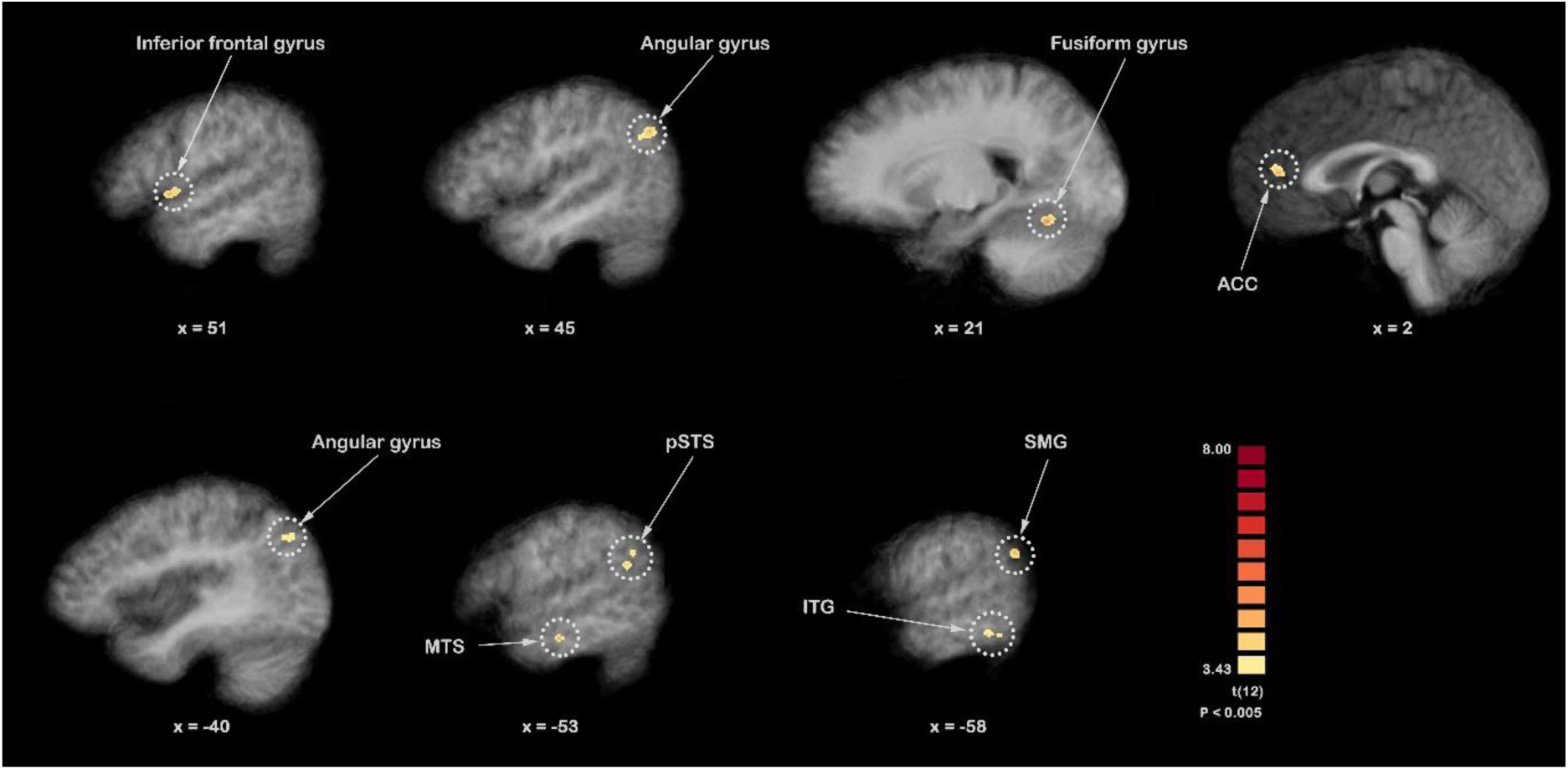
Clusters resulting from the fMRI GNB classification of emotion. The unsmoothed searchlight data of each individual participant served as input for the GNB classifier, separately. Subsequently, a group-level one-sample t-test against 0 was performed (2-tailed). The clusters are displayed at uncorrected p = 0.005 with a cluster threshold = 10 voxels. After correction for multiple comparisons (cluster size corrected with Monte-Carlo simulation, alpha level = 0.05, initial p = 0.005, numbers of iterations = 5000), no clusters survived. See Table SR7 in Supplementary Results for more details on their location and statistical values. Abbreviations: ACC: anterior cingulate cortex; ITG: inferior temporal gyrus; MTS: middle temporal sulcus; pSTS: posterior superior temporal sulcus; SMG: supramarginal gyrus.

The inferior frontal gyrus and precentral sulcus processed movements with comparable velocity values in a similar manner, regardless of the conveyed affect (see **Figure 5** and **Table SR5** in Supplementary Results). Previous research has implicated the inferior frontal gyrus in the observation of emotional actions (Dricu & Frühholz, 2016; Molenberghs, Cunnington, & Mattingley, 2012) and the precentral sulcus in motor preparation and execution (Grèzes, Armony, Rowe, & Passingham, 2003). The involvement of these areas in the processing of body movements is also in line with the hypothesis that observing body expressions triggers action preparation processes (Caspers et al., 2010; de Gelder et al., 2004; Fadiga, Craighero, & Olivier, 2005; Grafton & Hamilton, 2007). The current results suggest that the velocity of body movements may be used in these areas for the preparation of an appropriate behavioural response.

In general, more overlap was found among clusters correlating to postural features than to kinematic cues. Specifically, shoulder ratio, surface and limb contraction showed a high amount of overlap in subcortical areas, including caudate nucleus, putamen, insula and internal capsule (see **Figure 5** and **Table SR5** in Supplementary Results). However, the stimuli representation in these areas only presented positive correlations to limb contraction. This feature showed clear differences between fear and the rest of the stimuli categories and was negatively correlated to the features of shoulder ratio and surface, although not significantly (see **Figure 2** and **Table SR1** in Supplementary Results). In these latter features, both neutral and fearful conditions were represented differently from angry and, especially, happy expressions. Taking all this together, these findings suggest that the stimuli representation in these overlapping clusters is more similar to the representation reflected by limb contraction.

Limb angles showed positive correlations to several areas that have previously been implicated in the processing of emotions: precentral gyrus, anterior insula, superior temporal sulcus and postcentral gyrus (Dricu & Frühholz, 2016; Kober et al., 2008). Particularly, these regions have been suggested to play a role in action observation and motor preparation (i.e. precentral and poscentral gyrus; Caspers et al., 2010; Grafton & Hamilton, 2007; Valchev, Gazzola, Avenanti, & Keysers, 2016), interoception (i.e. anterior insula; Craig, 2009; Critchley, 2005; Karnath, Baier, & Nägele, 2005) and biological motion processing (i.e. superior temporal sulcus; Allison et al., 2000; Grossman et al., 2000; Grossman et al., 2010). Their similarity to the stimuli representation of limb angles indicates that these areas may be relevant in the distinction between neutral and emotional categories, as well as between fearful expressions and the rest of the movement categories (see **Figure 1**).

### Limb contraction and fear perception

Limb contraction not only showed positive correlations to subcortical structures but also to cortical areas. The observed clusters spanned affective, action observation, and motor preparation and execution networks, also known to be involved in the processing of emotional body expressions, especially fearful ones (de Gelder, 2006). In this regard, one of the most well-known areas implicated in the decoding of affective information from sensory relevant inputs (LeDoux, 2003), the amygdala, showed a positive correlation to the stimuli representation of limb contraction. For many decades, this subcortical area has been suggested to have a central role in the rapid perception and response to fear (Hadjikhani & de Gelder, 2003). Through its connections to sensory cortical regions, this area is thought to modulate attentional and perceptual processes and tune the motor system to initiate adaptive behaviours (Emery & Amaral, 2000). As the amygdala, another area that has been involved in the emotional labelling of incoming visual signals and that also positively correlated to limb contraction was the temporal pole (Olson, Plotzker, & Ezzyat, 2007).

Limb contraction also positively correlated to areas known to be implicated in emotional regulation and body awareness. One of these areas is the insula, a cortical region that has been suggested to integrate information about the location and condition of our bodies, our subjective emotions and the key features of our environment. Thus, it is believed that this area is key in associating internal and external experiences (Craig, 2009; Critchley, 2005; Karnath et al., 2005). Other areas also thought to play a role in monitoring the internal emotional state as well as in attention selection and planning, correlated positively to limb contraction, including the ACC (Devinsky, Morrell, & Vogt, 1995), the orbitofrontal cortex (Beer, John, Scabini, & Knight, 2006) and the dorsolateral prefrontal cortex (Phillips, Drevets, Rauch, & Lane, 2003).

As previously mentioned, limb contraction positively correlated to areas known to be involved in action understanding and motor preparation. One of these areas is the caudate nucleus, a subcortical region that has been implicated in the automatic and rapid perception of emotional bodies (de Gelder, 2006) as well as in goal-directed behaviours, by integrating information related to motor behaviour, actions and space (Grahn, Parkinson, & Owen, 2008). Particularly, the caudate nucleus may influence motor planning due to its connections with motor cortices (Utter & Basso, 2008) and structures that have been implicated in the affective evaluation of the environment, such as the insula (Chikama, McFarland, Amaral, & Haber, 1997; Craig, 2009) and the amygdala (Emery & Amaral, 2000). Interestingly, these areas were also positively correlated to limb contraction, as mentioned above. Another subcortical area that has shown involvement in motor planning and execution, the putamen (Grahn et al., 2008), also correlated positively to limb contraction. Cortical areas also suggested to be part of the action observation network and thought to be involved in motor preparation were observed, including pIPS, PMv, pre-SMA and IPL. Particularly, pIPS has been implicated in attention and action observation (Caspers et al., 2010; Corbetta et al., 2008; Culham & Valyear, 2006; Grafton & Hamilton, 2007) while PMv and pre-SMA have been involved in space perception, action and intention understanding (Lau et al., 2004; Rizzolatti et al., 1996; Rizzolatti et al., 2002; Urgesi, Calvo-Merino, et al., 2007; Urgesi, Candidi, et al., 2007), as well as in action preparation and execution (Hoshi & Tanji, 2007; Isoda, 2005; Luppino et al., 1991; Picard & Strick, 2001).

In summary, these findings underscore the important role of postural cues in the understanding of emotion from other people’s movements (Poyo Solanas et al., 2020). In particular, it seems that the contraction of the limbs might be the low-level visual property that drives the discrimination of fearful bodies across subcortical and cortical areas. In fact, the clusters showing a positive correlation to this feature are known to be involved in the processing of emotional body expressions, especially fearful ones (de Gelder, 2006; Meeren, Hadjikhani, Ahlfors, Hämäläinen, & De Gelder, 2016). These areas spanned affective, action observation and motor preparation networks, suggesting that in the course of body movement perception, the perceived affect and intent need to be integrated with different internal body signals in order to select and prepare for an appropriate and quick behavioural response. Moreover, the results suggest that the amygdala may be at the core of all these areas, since this structure is known to be crucial for the processing of fearful body expressions (Hadjikhani & de Gelder, 2003) and presents connections to many of the observed brain regions (Emery & Amaral, 2000). Specifically, the amygdala is thought to modulate attentional and perceptual processes via its connections to sensory cortical regions and tune the motor system to initiate adaptive behaviours (Emery & Amaral, 2000).

### Distributed representation of emotion in the brain

The current study investigated the representation of emotion using different approaches (i.e. ROI RSA, whole-brain searchlight RSA and whole-brain GNB classification), which gave diverse results. Several of the pre-defined body-selective regions showed a non-significant but positive correlation to affective categories, including frontal, premotor, parietal and temporo-occipital regions (see **Figure 3**). The only significant positive correlation was found in SMA, an area that has been related to action representation and motor preparation (Nachev, Kennard, & Husain, 2008). Interestingly, it has previously been shown that this area is strongly modulated by emotion (Engelen et al., 2018; Oliveri et al., 2003; Rodigari & Oliveri, 2014). Based on its connections to the amygdala (Grèzes, Valabrègue, Gholipour, & Chevallier, 2014), it has been proposed that SMA may play a role in transforming affective experience into motor actions (Oliveri et al., 2003). However, further research is needed to understand the mechanisms underlying this transformation. In the current study, supplementary motor clusters resulted from a positive correlation to the feature of limb contraction. This suggests that SMA may use the contraction of the body limbs to discriminate the affect conveyed by body movements. Particularly, the stimuli representation of limb contraction indicates that SMA may be important in the discrimination of fearful body expressions (see **Figure 1**).

Whole-brain RSA using a multivariate approach revealed that the emotional content conveyed by body movements (i.e. emotion RDM) is coded in middle temporal and occipital areas (see **Figure 5** and **Table SR5** in Supplementary Results). The location of these areas differed from the ones of the pre-defined clusters. The region located in the middle temporal gyrus has been suggested to play a role in motion observation (Rizzolatti et al., 1996) as well as in the attribution of intentions to others (Brunet, Sarfati, Hardy-Baylé, & Decety, 2000). Further research needs to be conducted to understand the mechanisms underlying the processing of affect in these regions.

Finally, the representation of affect from body movements was also investigated using a GNB algorithm (Ontivero-Ortega et al., 2017). Although the resulting clusters do not overlap with the areas obtained with the aforementioned approaches, these regions are in agreement with previous findings showing some level of involvement in emotional processing: angular gyrus, fusiform gyrus, ACC, middle temporal sulcus, SMG and inferior temporal gyrus (Dricu & Frühholz, 2016; Kober et al., 2008) (see **Figure 6** and **Table SR7** in Supplementary Results). Two of these areas, the angular gyrus and the ACC, positively correlated to limb contraction (see **Figure 5** and **Table SR5** in Supplementary Results), indicating that these regions may use this body feature to discriminate between affective movements, especially fear. The SMG showed a negative correlation to acceleration, suggesting that this area may distinguish between different affective movements despite presenting similar acceleration values. Fusiform negatively correlated to shoulder ratio and surface, indicating that this region may process body movements in a way that does not correspond to their surface properties. This was also the case for the middle temporal sulcus, which not only showed negative correlation to surface, but also to kinematic information. Nevertheless, further research is needed to understand how these regions encode affective information and whether they use kinematic and postural features for this purpose.

### Addressing outstanding questions

Our results are relevant for two outstanding questions in the literature. First, studies investigating the representation of the face and the body have identified specialised areas and time windows (Bentin, Allison, Puce, Perez, & McCarthy, 1996; Downing et al., 2001; Peelen & Downing, 2005; Schwarzlose et al., 2005; Stekelenburg & de Gelder, 2004). Yet, when taking into account the affective information conveyed by bodies or faces, areas beyond the category-specific ones have systematically emerged in fMRI studies (Dricu & Frühholz, 2016; Kober et al., 2008), and specific activity patterns have been observed before the N170 window (Pizzagalli et al., 2002; van Heijnsbergen, Meeren, Grezes, & de Gelder, 2007). Our results suggest that this earlier activity may be related to the coding of specific features and may occur relatively independently of subsequent stimulus categorisation (van Heijnsbergen et al., 2007). A similar explanation may apply for activations outside the face and body areas found in fMRI studies of body expression perception (Goldberg, Preminger, & Malach, 2014; Grèzes et al., 2007). Secondly, the notion that specific features carry discriminating information may provide a better understanding of some puzzling neuropsychological findings. For example, many reports in the literature show that patients with focal damage as in prosopagnosia may still have intact recognition of the emotional expression while their category-specific perception of the face (or the body) is impaired (de Gelder, Vroomen, Pourtois, & Weiskrantz, 1999; Tranel, Damasio, & Damasio, 1988). The possibility of feature-based expression recognition provides a promising candidate to address that puzzle.

## Conclusions

The current findings show that body movements differentially activated brain regions based on their postural and kinematic characteristics and thus, these aspects might be encoded in these regions. Among these features, the degree of limb contraction seems to be particularly relevant for distinguishing fear from other affective movements. This feature was represented in several regions spanning affective, action observation and motor preparation networks. Our approach goes beyond classical methods of categorically mapping cognitive categories to brain activation/deactivation and instead attempts to find a brain basis for affective body and action perception, looking for movement features and how they are encoded in the brain.

## Acknowledgements

This work was supported by the European Research Council (ERC) FP7-IDEAS-ERC (Grant agreement number 295673; Emobodies), by the Future and Emerging Technologies (FET) Proactive Programme H2020-EU.1.2.2 (Grant agreement 824160; EnTimeMent) and by the Industrial Leadership Programme H2020-EU.1.2.2 (Grant agreement 825079; MindSpaces).

