## Supplementary material for "Limb contraction drives fear perception": SR

### Supplementary Materials

#### Region of Interest definition

The functional data of the localizer were used to identify several regions of interest (ROI) with regard to body perception. For this purpose, a fixed-effects whole-brain general linear model (GLM) was fitted to the 3-mm smoothed data of each participant individually. The generated regression models consisted the %-signal-transformed predictors of each stimulus category (i.e. body alone, face alone, voice alone, body-voice, face-voice) convolved with a two-gamma HRF and the z-transformed motion parameters as predictors of no interest. The considered ROIs include occipito-temporal areas that have previously shown a certain level of body specificity (three ROIs: fusiform body area (FBA), extrastriate body area (EBA), posterior superior temporal sulcus (pSTS)) (Downing, Jiang, Shuman, & Kanwisher, 2001; Kontaris, Wiggett, & Downing, 2009; Peelen & Downing, 2005; Schwarzlose, Baker, & Kanwisher, 2005; Vangeneugden, Peelen, Tadin, & Battelli, 2014), parietal and temporal areas thought to be implicated in attention and action observation (six ROIs: V7/3a, superior parieto-occipital cortex (SPOC), superior marginal gyrus (SMG), posterior intraparietal sulcus (pIPS), medial intraparietal sulcus (mIPS), anterior intraparietal sulcus (aIPS)) (Caspers, Zilles, Laird, & Eickhoff, 2010; Corbetta, Patel, & Shulman, 2008; Culham & Valyear, 2006; Grafton & Hamilton, 2007), and frontal areas involved in action observation and other higher cognitive functions (six ROIs: ventral and dorsal premotor cortex (PMv, PMd), supplementary motor area (SMA), pre-supplementary motor area (pre-SMA), inferior frontal and frontal regions) (Caspers et al., 2010; Grafton & Hamilton, 2007).

Area V3a/V7 was located on the IPS at the level of the occipito-parietal junction. SPOC was defined at the superior part of the occipito-parietal junction. SMG was defined in the ascending posterior ramus of the lateral fissure. pSTS was located at the posterior portion of the superior temporal sulcus. The IPS was divided into anterior, middle and posterior areas giving aIPS, mIPS, pIPS, respectively. PMv was defined at the intersection between the inferior frontal sulcus and the inferior precentral sulcus, whereas PMd at the junction between the superior frontal sulcus and the precentral sulcus. IFG was located in the inferior frontal sulcus, anterior to the defined PMv. Pre-SMA and SMA were defined on the medial surface of the superior frontal gyrus. EBA and FBA were defined as the clusters localized at the lateral occipital temporal lobe and fusiform gyri, respectively. All body movement videos were contrasted to baseline at uncorrected p = 0.005 to define all the ROIs with the exception of FBA and EBA. For these regions, body movements were contrasted against facial movement clips at uncorrected p = 0.005. The ROIs were defined bilaterally whenever possible and subsequently merged into a single ROI for each participant individually.

**Table SM1. ROI details**

| ROI | Average TAL coordinates | | | | | | Voxels | | |
|---|---|---|---|---|---|---|---|---|---|
| | X | X (SD) | Y | Y (SD) | Z | Z (SD) | Voxels (avg.) | Voxels (SD) | Number of participants |
| Frontal | -7.15 | 14.63 | 47.17 | 2.36 | 20.55 | 1.72 | 187 | 96.74 | 4 |
| IF | 20.34 | 19.70 | 26.42 | 3.54 | 16.39 | 3.14 | 502 | 163.14 | 9 |
| PMd | 15.71 | 21.68 | -5.10 | 2.30 | 47.34 | 3.35 | 1087 | 265.39 | 12 |
| PMv | 24.86 | 24.21 | 4.38 | 3.05 | 29.11 | 2.75 | 756 | 223.58 | 11 |
| PreSMA | 2.00 | 3.05 | 9.76 | 1.34 | 50.51 | 1.67 | 162 | 57.03 | 5 |
| SMA | 2.76 | 5.03 | -3.69 | 2.16 | 59.71 | 3.40 | 560 | 243.72 | 7 |
| aIPS | 7.49 | 16.74 | -47.20 | 3.95 | 47.33 | 3.03 | 910 | 267.07 | 8 |
| mIPS | 11.95 | 17.03 | -58.07 | 4.12 | 45.53 | 3.69 | 984 | 116.98 | 7 |
| pIPS | -4.78 | 14.48 | -72.27 | 3.33 | 26.29 | 3.31 | 842 | 271.40 | 11 |
| V73a | 1.28 | 13.27 | -86.56 | 3.17 | 21.03 | 3.66 | 1131 | 224.90 | 12 |
| SPOC | 4.94 | 9.82 | -73.13 | 2.43 | 32.66 | 2.50 | 434 | 162.86 | 12 |
| SMG | 34.79 | 28.34 | -37.41 | 3.54 | 22.34 | 3.22 | 1026 | 188.75 | 11 |
| pSTS | 11.79 | 39.95 | -43.94 | 4.32 | 11.05 | 3.61 | 1838 | 353.08 | 12 |
| EBA | 4.99 | 44.68 | -71.91 | 4.46 | 6.13 | 4.45 | 2603 | 360.22 | 13 |
| FBA | 11.25 | 23.57 | -56.33 | 3.55 | -11.11 | 2.64 | 651 | 200.06 | 9 |
| EVC | 0.14 | 8.60 | -91.96 | 3.19 | -6.01 | 3.97 | 1729 | 236.81 | 13 |

**Abbreviation:** AVG.: average; EBA: extrastriate body area; EVC: early visual cortex; FBA: fusiform body area; IF: inferior frontal cortex; IPS: intraparietal sulcus, p: posterior, m: middle, a: anterior; PMd: dorsal premotor cortex; PMv: ventral premotor cortex; pre-SMA: pre-supplementary motor area; pSTS: posterior superior temporal sulcus; SD: standard deviation; SMA: supplementary motor area; SMG: supramarginal gyrus; SPOC: superior parietal occipital cortex.

**Univariate analyses**

In order to check the quality of the data, a fixed-effects whole-brain GLM was fitted to the unsmoothed data of the main experiment for each participant individually. Also, a random-effects whole-brain GLM was performed at the group level with the 4mm-smoothed functional data. The generated regression models consisted the %-signal-transformed predictors of each stimulus category (i.e. angry bodies, happy bodies, neutral bodies and fearful bodies) convolved with a two-gamma HRF and the z-transformed motion parameters as predictors of no interest (see Figure SR2 in Supplementary Results to visualize the resulting clusters at the group level).

### Supplementary Results

**TABLE SR1. Correlation between representational dissimilarity matrices of kinematic and postural features.** The correlations were computed using Spearman's rank correlation. Both uncorrected and Bonferroni-corrected p-values are displayed for each comparison ($\alpha_{bonf}$ = 0.05/9, with nine comparisons per feature).

| | Emotional categories | | | Velocity | | | Acceleration | | | Vertical movement | | |
|---|---|---|---|---|---|---|---|---|---|---|---|---|
| | r | p-value | corr. p-value | r | p-value | corr. p-value | r | p-value | corr. p-value | r | p-value | corr. p-value |
| **Velocity** | 0.090 | 0.330 | 0.999 | | | | | | | | | |
| **Acceleration** | 0.153 | 0.096 | 0.862 | 0.712 | 0.000 | 0.000 | | | | | | |
| **Vertical movement** | 0.038 | 0.676 | 0.999 | 0.369 | 0.000 | 0.000 | 0.087 | 0.342 | 0.999 | | | |
| **Limb angles** | 0.408 | 0.000 | 0.000 | -0.054 | 0.554 | 0.999 | 0.045 | 0.624 | 0.999 | -0.304 | 0.001 | 0.007 |
| **Symmetry** | 0.349 | 0.000 | 0.001 | -0.097 | 0.289 | 0.999 | -0.004 | 0.963 | 0.999 | -0.266 | 0.003 | 0.031 |
| **Shoulder ratio** | 0.345 | 0.000 | 0.001 | 0.205 | 0.025 | 0.226 | 0.001 | 0.990 | 0.999 | 0.124 | 0.176 | 0.999 |
| **Surface** | 0.215 | 0.018 | 0.164 | 0.216 | 0.018 | 0.162 | 0.004 | 0.964 | 0.999 | 0.281 | 0.002 | 0.018 |
| **Limb contraction** | 0.064 | 0.485 | 0.999 | -0.180 | 0.050 | 0.447 | 0.061 | 0.509 | 0.999 | -0.202 | 0.027 | 0.245 |

**Abbreviation**: Bonf.: Bonferroni; Corr.: corrected.

**TABLE SR1 (continuation). Correlation between representational dissimilarity matrices of kinematic and postural features.** The correlations were computed using Spearman's rank correlation. Both uncorrected and Bonferroni-corrected p-values are displayed for each comparison ($\alpha_{bonf}$ = 0.05/9, with nine comparisons per feature).

| | **Limb angles** | | | **Symmetry** | | | **Shoulder ratio** | | | **Surface** | | |
|---|---|---|---|---|---|---|---|---|---|---|---|---|
| | r | p-value | corr. p-value | r | p-value | corr. p-value | r | p-value | corr. p-value | r | p-value | corr. p-value |
| **Symmetry** | 0.555 | 0.000 | 0.000 | | | | | | | | | |
| **Shoulder ratio** | 0.450 | 0.000 | 0.000 | 0.323 | 0.000 | 0.003 | | | | | | |
| **Surface** | 0.291 | 0.001 | 0.012 | 0.007 | 0.940 | 0.999 | 0.741 | 0.000 | 0.000 | | | |
| **Limb contraction** | 0.489 | 0.000 | 0.000 | 0.067 | 0.463 | 0.999 | -0.169 | 0.066 | 0.590 | -0.053 | 0.564 | 0.999 |

**Abbreviation**: Bonf.: Bonferroni; Corr.: corrected.

**Table SR2. Results of pair-sample t-tests comparing kinematic and postural feature processing in each individual ROI.** The kinematic features consisted of velocity, acceleration and vertical movement. The postural features comprised shoulder ratio, surface, limb distances, symmetry and limb angles. Both uncorrected and Bonferroni-corrected p-values are displayed for each comparison ($\alpha_{bonf}$ = 0.05/16, with 16 comparisons).

| | **Kinematic features (Avg.)** | **Kinematic features (SD)** | **Postural features (Avg.)** | **Postural features (SD)** | **t-value** | **p-value** | **Corr. p-value** | **df** |
|---|---|---|---|---|---|---|---|---|
| **Frontal** | -0.04 | 0.07 | 0.07 | 0.08 | -1.42 | 0.251 | 0.999 | 3 |
| **IF** | -0.02 | 0.11 | 0.00 | 0.06 | -0.42 | 0.686 | 0.999 | 8 |
| **PMd** | 0.05 | 0.14 | 0.00 | 0.05 | 1.25 | 0.239 | 0.999 | 11 |
| **PMv** | -0.09 | 0.08 | 0.04 | 0.05 | -3.80 | 0.004 | 0.056 | 10 |
| **Pre-SMA** | 0.03 | 0.06 | 0.07 | 0.04 | -0.79 | 0.476 | 0.999 | 4 |
| **SMA** | -0.05 | 0.11 | 0.03 | 0.02 | -1.74 | 0.132 | 0.999 | 6 |
| **aIPS** | -0.06 | 0.08 | 0.02 | 0.08 | -1.60 | 0.154 | 0.999 | 7 |
| **mIPS** | 0.05 | 0.23 | 0.00 | 0.09 | 0.38 | 0.717 | 0.999 | 6 |
| **pIPS** | 0.00 | 0.14 | 0.00 | 0.09 | 0.03 | 0.978 | 0.999 | 10 |
| **V7** | 0.02 | 0.16 | 0.01 | 0.10 | 0.16 | 0.877 | 0.999 | 11 |
| **SPOC** | 0.00 | 0.12 | 0.01 | 0.07 | -0.32 | 0.756 | 0.999 | 11 |
| **SMG** | 0.00 | 0.13 | 0.00 | 0.05 | -0.04 | 0.969 | 0.999 | 10 |
| **pSTS** | -0.02 | 0.10 | 0.04 | 0.07 | -1.40 | 0.190 | 0.999 | 11 |
| **EBA** | -0.07 | 0.13 | 0.03 | 0.10 | -1.76 | 0.104 | 0.999 | 12 |
| **FBA** | -0.06 | 0.10 | 0.01 | 0.05 | -1.39 | 0.203 | 0.999 | 8 |
| **EVC** | -0.06 | 0.12 | 0.03 | 0.06 | -1.76 | 0.104 | 0.999 | 12 |

**Abbreviation:** Avg.: average; corr.: corrected; df: degrees of freedom; EBA: extrastriate body area; EVC: early visual cortex; FBA: fusiform body area; IF: inferior frontal cortex; IPS: intraparietal sulcus, p: posterior, m: middle, a: anterior; PMd: dorsal premotor cortex; PMv: ventral premotor cortex; pre-SMA: pre-supplementary motor area; pSTS: posterior superior temporal sulcus; SD: standard deviation; SMA: supplementary motor area; SMG: supramarginal gyrus; SPOC: superior parietal occipital cortex.

**Table SR3. Results of pair-sample t-tests comparing the processing of each feature in dorsal and ventral clusters.** The dorsal clusters consisted of aIPS, mIPS, pIPS, V7/3a, SPOC, SMG and pSTS. The ventral clusters consisted of EBA and FBA. Both uncorrected and Bonferroni-corrected p-values are displayed for each comparison ($\alpha_{bonf}$ = 0.05/9, with nine comparisons).

| | **Dorsal clusters (Avg.)** | **Dorsal clusters (SD)** | **Ventral clusters (Avg.)** | **Ventral clusters (SD)** | **t-value** | **p-value** | **Corr. p-value** | **df** |
|---|---|---|---|---|---|---|---|---|
| **Emotional category** | 0.01 | 0.03 | -0.01 | 0.04 | 0.86 | 0.408 | 0.999 | 12 |
| **Velocity** | 0.03 | 0.15 | -0.06 | 0.15 | 2.23 | 0.046 | 0.411 | 12 |
| **Acceleration** | 0.00 | 0.11 | -0.04 | 0.14 | 1.26 | 0.230 | 0.999 | 12 |
| **Vertical movement** | 0.00 | 0.15 | -0.06 | 0.16 | 1.14 | 0.276 | 0.999 | 12 |
| **Shoulder ratio** | -0.01 | 0.05 | -0.01 | 0.05 | 0.35 | 0.729 | 0.999 | 12 |
| **Surface** | -0.01 | 0.07 | -0.05 | 0.09 | 1.43 | 0.177 | 0.999 | 12 |
| **Limb contraction** | 0.04 | 0.12 | 0.08 | 0.14 | -1.17 | 0.263 | 0.999 | 12 |
| **Symmetry** | -0.01 | 0.09 | 0.01 | 0.11 | -0.81 | 0.435 | 0.999 | 12 |
| **Limb angles** | 0.02 | 0.14 | 0.05 | 0.14 | -1.16 | 0.270 | 0.999 | 12 |

**Abbreviation:** Avg.: average; Corr.: corrected; EBA: extrastriate body area; df: degrees of freedom; FBA: fusiform body area; IPS: intraparietal sulcus, p: posterior, m: middle, a: anterior; pSTS: posterior superior temporal sulcus; SD: standard deviation; SMG: supramarginal gyrus; SPOC: superior parietal occipital cortex.

**Table SR4. Correlation between ROI representational dissimilarity matrices**. The correlations were computed using Spearman's rank correlation. ROI RDMs were performed at the participant level first, and the resulting matrix was then averaged across participants. Both corrected and Bonferroni-corrected p-values are displayed for each comparison ($\alpha_{bonf}$ = 0.05/16, with 16 comparisons per ROI).

| | **Frontal** | | | **IF** | | | **PMd** | | | **PMv** | | |
|---|---|---|---|---|---|---|---|---|---|---|---|---|
| | r-value | p-value | corr. p-value | r-value | p-value | corr. p-value | r-value | p-value | corr. p-value | r-value | p-value | corr. p-value |
| **IF** | 0.00 | 0.992 | 0.999 | | | | | | | | | |
| **PMd** | 0.02 | 0.842 | 0.999 | 0.19 | 0.039 | 0.624 | | | | | | |
| **PMv** | 0.10 | 0.286 | 0.999 | -0.08 | 0.398 | 0.999 | -0.20 | 0.025 | 0.401 | | | |
| **Pre-SMA** | -0.18 | 0.055 | 0.884 | -0.08 | 0.359 | 0.999 | 0.12 | 0.194 | 0.999 | 0.03 | 0.740 | 0.999 |
| **SMA** | -0.03 | 0.775 | 0.999 | 0.07 | 0.425 | 0.999 | 0.03 | 0.727 | 0.999 | 0.22 | 0.015 | 0.245 |
| **aIPS** | 0.05 | 0.582 | 0.999 | 0.09 | 0.307 | 0.999 | 0.20 | 0.029 | 0.458 | 0.18 | 0.045 | 0.724 |
| **mIPS** | 0.12 | 0.199 | 0.999 | 0.22 | 0.017 | 0.277 | 0.43 | 0.000 | 0.000 | 0.01 | 0.892 | 0.999 |
| **pIPS** | -0.19 | 0.038 | 0.608 | 0.34 | 0.000 | 0.002 | 0.28 | 0.002 | 0.027 | -0.01 | 0.921 | 0.999 |
| **V7/3a** | -0.12 | 0.182 | 0.999 | 0.14 | 0.133 | 0.999 | 0.28 | 0.002 | 0.028 | -0.06 | 0.511 | 0.999 |
| **SPOC** | -0.08 | 0.391 | 0.999 | 0.01 | 0.877 | 0.999 | 0.08 | 0.364 | 0.999 | 0.35 | 0.000 | 0.001 |
| **SMG** | -0.07 | 0.472 | 0.999 | 0.40 | 0.000 | 0.000 | 0.11 | 0.252 | 0.999 | 0.20 | 0.026 | 0.414 |
| **pSTS** | 0.11 | 0.219 | 0.999 | 0.36 | 0.000 | 0.001 | 0.36 | 0.000 | 0.001 | 0.31 | 0.001 | 0.009 |
| **EBA** | 0.01 | 0.887 | 0.999 | 0.41 | 0.000 | 0.000 | 0.23 | 0.011 | 0.178 | 0.13 | 0.172 | 0.999 |
| **FBA** | 0.11 | 0.232 | 0.999 | 0.00 | 0.964 | 0.999 | -0.15 | 0.105 | 0.999 | 0.20 | 0.032 | 0.518 |
| **EVC** | 0.15 | 0.091 | 0.999 | -0.01 | 0.914 | 0.999 | 0.03 | 0.715 | 0.999 | 0.19 | 0.042 | 0.664 |

**Abbreviation**: Bonf.: Bonferroni; Corr.: corrected; EBA: extrastriate body area; EVC: early visual cortex; FBA: fusiform body area; IF: inferior frontal cortex; IPS: intraparietal sulcus, p: posterior, m: middle, a: anterior; PMd: dorsal premotor cortex; PMv: ventral premotor cortex; pre-SMA: pre-supplementary motor area; pSTS: posterior superior temporal sulcus; SD: standard deviation; SMA: supplementary motor area; SMG: supramarginal gyrus; SPOC: superior parietal occipital cortex.

**Table SR4 (continuation). Correlation between ROI representational dissimilarity matrices**. The correlations were computed using Spearman's rank correlation. ROI RDMs were performed at the participant level first, and the resulting matrix was then averaged across participants. Both corrected and Bonferroni-corrected p-values are displayed for each comparison ($\alpha_{bonf}$ = 0.05/16, with 16 comparisons per ROI).

| | **preSMA** | | | **SMA** | | | **IPSa** | | | **IPSm** | | |
|---|---|---|---|---|---|---|---|---|---|---|---|---|
| | r-value | p-value | corr. p-value | r-value | p-value | corr. p-value | r-value | p-value | corr. p-value | r-value | p-value | corr. p-value |
| **SMA** | 0.06 | 0.500 | 0.999 | | | | | | | | | |
| **aIPS** | -0.13 | 0.166 | 0.999 | 0.08 | 0.410 | 0.999 | | | | | | |
| **mIPS** | -0.02 | 0.827 | 0.999 | -0.02 | 0.815 | 0.999 | 0.41 | 0.000 | 0.000 | | | |
| **pIPS** | -0.11 | 0.231 | 0.999 | -0.08 | 0.364 | 0.999 | 0.23 | 0.012 | 0.194 | 0.19 | 0.035 | 0.566 |
| **V7/3a** | 0.02 | 0.803 | 0.999 | 0.08 | 0.381 | 0.999 | 0.26 | 0.004 | 0.060 | 0.22 | 0.016 | 0.260 |
| **SPOC** | 0.15 | 0.103 | 0.999 | 0.13 | 0.142 | 0.999 | 0.34 | 0.000 | 0.003 | 0.28 | 0.002 | 0.031 |
| **SMG** | -0.07 | 0.462 | 0.999 | 0.11 | 0.229 | 0.999 | 0.05 | 0.579 | 0.999 | 0.20 | 0.025 | 0.407 |
| **pSTS** | 0.15 | 0.100 | 0.999 | 0.29 | 0.001 | 0.020 | 0.21 | 0.022 | 0.352 | 0.32 | 0.000 | 0.007 |
| **EBA** | -0.11 | 0.232 | 0.999 | 0.13 | 0.153 | 0.999 | 0.21 | 0.022 | 0.346 | 0.15 | 0.094 | 0.999 |
| **FBA** | 0.16 | 0.074 | 0.999 | 0.19 | 0.037 | 0.587 | 0.09 | 0.336 | 0.999 | -0.16 | 0.087 | 0.999 |
| **EVC** | 0.17 | 0.066 | 0.999 | 0.31 | 0.001 | 0.008 | 0.19 | 0.034 | 0.537 | 0.02 | 0.826 | 0.999 |

**Abbreviation**: Bonf.: Bonferroni; Corr.: corrected; EBA: extrastriate body area; EVC: early visual cortex; FBA: fusiform body area; IF: inferior frontal cortex; IPS: intraparietal sulcus, p: posterior, m: middle, a: anterior; PMd: dorsal premotor cortex; PMv: ventral premotor cortex; pre-SMA: pre-supplementary motor area; pSTS: posterior superior temporal sulcus; SD: standard deviation; SMA: supplementary motor area; SMG: supramarginal gyrus; SPOC: superior parietal occipital cortex.

**Table SR4 (continuation). Correlation between ROI representational dissimilarity matrices**. The correlations were computed using Spearman's rank correlation. ROI RDMs were performed at the participant level first, and the resulting matrix was then averaged across participants. Both corrected and Bonferroni-corrected p-values are displayed for each comparison ($\alpha_{bonf}$ = 0.05/16, with 16 comparisons per ROI).

| | **IPSp** | | | **V73a** | | | **SPOC** | | | **SMG** | | |
|---|---|---|---|---|---|---|---|---|---|---|---|---|
| | r-value | p-value | corr. p-value | r-value | p-value | corr. p-value | r-value | p-value | corr. p-value | r-value | p-value | corr. p-value |
| **V73a** | 0.37 | 0.000 | 0.001 | | | | | | | | | |
| **SPOC** | 0.18 | 0.054 | 0.863 | 0.44 | 0.000 | 0.000 | | | | | | |
| **SMG** | 0.47 | 0.000 | 0.000 | 0.44 | 0.000 | 0.000 | 0.34 | 0.000 | 0.003 | | | |
| **pSTS** | 0.33 | 0.000 | 0.005 | 0.20 | 0.031 | 0.498 | 0.33 | 0.000 | 0.005 | 0.58 | 0.000 | 0.000 |
| **EBA** | 0.57 | 0.000 | 0.000 | 0.33 | 0.000 | 0.004 | 0.11 | 0.242 | 0.999 | 0.64 | 0.000 | 0.000 |
| **FBA** | -0.24 | 0.009 | 0.148 | 0.28 | 0.002 | 0.030 | 0.20 | 0.029 | 0.465 | 0.05 | 0.562 | 0.999 |
| **EVC** | 0.00 | 0.994 | 0.999 | 0.28 | 0.002 | 0.034 | 0.30 | 0.001 | 0.012 | 0.03 | 0.715 | 0.999 |

**Abbreviation**: Bonf.: Bonferroni; Corr.: corrected; EBA: extrastriate body area; EVC: early visual cortex; FBA: fusiform body area; IF: inferior frontal cortex; IPS: intraparietal sulcus, p: posterior, m: middle, a: anterior; PMd: dorsal premotor cortex; PMv: ventral premotor cortex; pre-SMA: pre-supplementary motor area; pSTS: posterior superior temporal sulcus; SD: standard deviation; SMA: supplementary motor area; SMG: supramarginal gyrus; SPOC: superior parietal occipital cortex.

**Table SR4 (continuation). Correlation between ROI representational dissimilarity matrices**. The correlations were computed using Spearman's rank correlation. ROI RDMs were performed at the participant level first, and the resulting matrix was then averaged across participants. Both corrected and Bonferroni-corrected p-values are displayed for each comparison ($\alpha_{bonf}$ = 0.05/16, with 16 comparisons per ROI).

| | **pSTS** | | | **EBA** | | | **FBA** | | |
|---|---|---|---|---|---|---|---|---|---|
| | r-value | p-value | corr. p-value | r-value | p-value | corr. p-value | r-value | p-value | corr. p-value |
| **EBA** | 0.57 | 0.000 | 0.000 | | | | | | |
| **FBA** | 0.01 | 0.895 | 0.999 | -0.01 | 0.871 | 0.999 | | | |
| **EVC** | 0.30 | 0.001 | 0.013 | 0.08 | 0.361 | 0.999 | 0.34 | 0.000 | 0.002 |

**Abbreviation**: Bonf.: Bonferroni; Corr.: corrected; EBA: extrastriate body area; EVC: early visual cortex; FBA: fusiform body area; IF: inferior frontal cortex; IPS: intraparietal sulcus, p: posterior, m: middle, a: anterior; PMd: dorsal premotor cortex; PMv: ventral premotor cortex; pre-SMA: pre-supplementary motor area; pSTS: posterior superior temporal sulcus; SD: standard deviation; SMA: supplementary motor area; SMG: supramarginal gyrus; SPOC: superior parietal occipital cortex.

**Table SR5. Clusters resulting from the searchlight RSA of the features.** The searchlight RSA dissimilarity matrices were correlated with each of the feature RDMs, respectively. The resulting maps were z-transformed for each participant. Subsequently, group-level one-sample t-tests against 0 (2-tailed, cluster size corrected with Monte-Carlo simulation, alpha level = 0.05, initial p = 0.005, numbers of iterations = 5000) were performed, one for each feature.

| Feature | Location | H | TAL mean x | TAL mean y | TAL mean z | SD x | SD y | SD z | Cluster size ($mm^3$) | Avg. r-value | Avg. t-value | Avg. p-value |
|---|---|---|---|---|---|---|---|---|---|---|---|---|
| **Emotional categories** | MTG | R | 61.27 | -13.3 | -17.45 | 1.91 | 2.7 | 1.86 | 3128 | .068 | 3.967 | .002 |
| | MOG | L | -28.41 | -90.07 | -4.76 | 2.74 | 3.23 | 3.35 | 5960 | .061 | 4.082 | .002 |
| **Velocity** | IFG | R | 29.09 | 21.61 | 28.38 | 2.11 | 2.87 | 2.57 | 4176 | .089 | 3.987 | .002 |
| | Central sulcus | L | -27.54 | -22.26 | 37.13 | 5.41 | 3.48 | 5.17 | 13456 | .077 | 4.124 | .002 |
| | Angular gyrus | L | -43.19 | -75.43 | 30.58 | 3.59 | 3.05 | 2.26 | 4952 | -.061 | -4.251 | .002 |
| | MTG | L | -54.81 | -15.43 | -13.23 | 1.83 | 3.5 | 2.12 | 4040 | -.071 | -4.412 | .002 |
| **Acceleration** | MOG | R | 43.82 | -67.07 | 7.76 | 3.26 | 3.25 | 3.84 | 11200 | -.127 | -4.172 | .002 |
| | Ventricle | R | 21.02 | -32.31 | 21.81 | 2.26 | 2.57 | 2.44 | 3808 | -.067 | -4.153 | .002 |
| | Corpus callosum | R | 17.36 | 18.26 | 21.09 | 2.11 | 2.96 | 2.45 | 3984 | -.071 | -3.819 | .003 |
| | MTG (posterior part) | L | -47.48 | -61.32 | 1.49 | 6.04 | 4.17 | 4.1 | 25280 | -.122 | -4.068 | .002 |
| | SMG | L | -47.18 | -40.07 | 32.31 | 3.73 | 3.07 | 2.37 | 5320 | -.067 | -3.754 | .003 |
| | MTG | L | -49.52 | 1.36 | -21.38 | 4.11 | 3.53 | 3.73 | 12072 | -.075 | -4.294 | .002 |
| **Vertical movement** | STS (posterior part) | L | -44.64 | -41.72 | 3.56 | 2.21 | 2.99 | 2.34 | 4392 | -.091 | -4.006 | .002 |
| **Limb angles** | Precentral gyrus | R | 46.12 | -0.07 | 43.82 | 2.31 | 2.48 | 2.28 | 4312 | .072 | 3.870 | .003 |
| | Anterior insula | L | -29.72 | 16.1 | -10.86 | 3.55 | 2.2 | 4.24 | 6264 | .073 | 4.028 | .002 |
| | STS | L | -51.62 | -51.02 | 7.53 | 2.83 | 4.93 | 4.48 | 13648 | .095 | 4.012 | .002 |
| | Postcentral gyrus | L | -60.94 | -19.2 | 30.12 | 1.93 | 3.33 | 2.21 | 3640 | .055 | 3.949 | .002 |
| **Symmetry** | IOG | R | 30.53 | -81.2 | -19.53 | 2.87 | 3.15 | 2.13 | 3712 | -.054 | -3.859 | .003 |
| | Anterior calcarine sulcus, Isthmus | L | -12.98 | -47.05 | 4.72 | 3.51 | 3.51 | 3.15 | 10200 | -.076 | -3.939 | .002 |
| | Central insular sulcus | L | -23.91 | 10.52 | -20.39 | 2.13 | 2.8 | 2.82 | 4664 | .072 | 3.775 | .003 |
| **Shoulder ratio** | CN, IC, Putamen, Insula | R | 22.05 | 21.3 | 7.83 | 4.88 | 7.86 | 4.69 | 32352 | -.072 | -4.247 | .002 |
| | Putamen, Claustrum, Amygdala | R | 29.21 | -2.42 | -6.36 | 3.2 | 1.63 | 4 | 4488 | -.062 | -3.772 | .003 |
| | Hippocampus | R | 19.75 | -33.17 | -0.43 | 3.13 | 3.46 | 2.77 | 4112 | -.063 | -3.800 | .003 |
| | Corpus callosum | R | 5.69 | -7.91 | 24.22 | 3.77 | 2.17 | 2.04 | 3568 | -.080 | -3.753 | .003 |
| | Cingulate gyrus | L | -0.53 | -44.99 | 34.6 | 2.93 | 2.36 | 1.66 | 3648 | -.062 | -3.989 | .002 |
| | Isthmus, cerebellum | L | -5.32 | -42.98 | -3.76 | 3.76 | 5.24 | 4.63 | 11592 | -.058 | -3.984 | .002 |
| | SFS, IFS, Anterior insula, CN, subACC | L | -23.3 | 26.56 | 11.53 | 11.11 | 5.81 | 10.24 | 69256 | -.073 | -4.298 | .002 |
| | CN, Putamen, IC | L | -18.51 | 0.27 | 11.95 | 6.97 | 5.26 | 7.49 | 13496 | -.070 | -3.971 | .002 |
| | Parahippocampal gyrus | L | -32.43 | -43.08 | 0.04 | 3.43 | 4.04 | 4.44 | 8336 | -.071 | -4.053 | .002 |
| | Central insular sulcus | L | -36.37 | 0.56 | -17.76 | 1.97 | 4.16 | 2.19 | 3064 | -.069 | -3.676 | .003 |
| | Fusiform gyrus | L | -38.01 | -15.79 | -20.26 | 2.29 | 1.98 | 2.11 | 3096 | -.062 | -3.997 | .002 |

**Abbreviation:** ACC: anterior cingulate gyrus; Avg.: averaged; CN: caudate nucleus; H: hemisphere; IC: internal capsule; IFG: inferior frontal gyrus; IFS: inferior frontal sulcus; IOG: inferior occipital gyrus; IPL: inferior parietal lobe; L: left hemisphere; MFG: middle frontal gyrus; MOG: middle occipital gyrus; MTG: middle temporal gyrus; MTS: middle temporal sulcus; R: right hemisphere; SD: standard deviation; SFG: superior frontal gyrus; SFS: superior frontal sulcus; SMG: supramarginal gyrus; STS: superior temporal sulcus; SubACC: subgenual anterior cingulate cortex; TAL: Talairach.

**Table SR5 (continuation). Clusters resulting from the searchlight RSA of the features.** The searchlight RSA dissimilarity matrices were correlated with each of the feature RDMs, respectively. The resulting maps were z-transformed for each participant. Subsequently, group-level one-sample t-tests against 0 (2-tailed, cluster size corrected with Monte-Carlo simulation, alpha level = 0.05, initial p = 0.005, numbers of iterations = 5000) were performed, one for each feature.

| Feature | Location | H | TAL mean x | TAL mean y | TAL mean z | SD x | SD y | SD z | Cluster size ($mm^3$) | Avg. r-value | Avg. t-value | Avg. p-value |
|---|---|---|---|---|---|---|---|---|---|---|---|---|
| **Surface** | Anterior insula, CN | R | 20.24 | 24.48 | 1.88 | 4.21 | 3.79 | 3.7 | 15880 | -.077 | -4.277 | .002 |
| | Thalamus | R | 18.18 | -26.86 | 3.93 | 3.41 | 4.36 | 4.3 | 13640 | -.079 | -4.013 | .002 |
| | White matter | R | 21.27 | 8.4 | 32.88 | 2.31 | 2.92 | 3.28 | 5464 | -.056 | -3.932 | .002 |
| | SubACC, ACC, Anterior insula, Frontal operculum, SFS | L | -15.74 | 29.87 | 12.64 | 7.39 | 4.2 | 11.63 | 33408 | -.075 | -4.148 | .002 |
| | CN, IC, Putamen | L | -18.69 | -6.84 | 17.54 | 3.75 | 3.81 | 2.9 | 7232 | -.064 | -3.929 | .002 |
| | Fusiform gyrus | L | -29.06 | -73.08 | -6.68 | 5.78 | 2.02 | 2.02 | 4552 | -.064 | -4.048 | .002 |
| | MTS | L | -42.29 | -34.74 | -6.37 | 6.74 | 5.5 | 4.48 | 6640 | -.071 | -3.905 | .003 |
| **Limb contraction** | Precentral sulcus (inferior part) | R | 47.39 | -1.65 | 42.88 | 2.3 | 2.72 | 4.19 | 5168 | .086 | 3.746 | .003 |
| | IPL (posterior part) | R | 31.2 | -69.12 | 28.23 | 3.6 | 5.21 | 4.13 | 10336 | .133 | 3.924 | .002 |
| | Insula, Putamen, Amygdala | R | 27.53 | -3.9 | -0.39 | 4.22 | 2.8 | 9.7 | 14992 | .081 | 3.953 | .002 |
| | Anterior insula, CN | R | 19.36 | 18.7 | 6.64 | 9.22 | 6.1 | 4.79 | 19552 | .092 | 3.940 | .002 |
| | SFG | R | 15.35 | -6.78 | 57.68 | 4.26 | 8.25 | 2.34 | 14352 | .083 | 3.974 | .002 |
| | SFG | R | 8.15 | 17.98 | 53.45 | 4.08 | 3.13 | 2.76 | 8328 | .088 | 4.232 | .002 |
| | Corpus callosum | R | 6.91 | -19.54 | 25.46 | 3.13 | 3.2 | 2.13 | 5600 | .091 | 4.031 | .002 |
| | ACC, anterior orbital gyrus | L | -17.29 | 36.36 | 5.75 | 4.52 | 3.4 | 7.12 | 15208 | .087 | 3.918 | .003 |
| | Anterior insula | L | -26.02 | 21.19 | 8.67 | 4.92 | 3.61 | 3.37 | 10600 | .066 | 4.352 | .002 |
| | MFG | L | -28.51 | 32.7 | 37.44 | 2 | 4.28 | 2.58 | 5344 | .070 | 3.884 | .003 |
| | Precentral sulcus (inferior part)/ central sulcus | L | -42.31 | -8.82 | 34.92 | 7.95 | 5.36 | 3.45 | 9200 | .072 | 3.840 | .003 |
| | Angular gyrus | L | -53.57 | -60.1 | 28.61 | 3.51 | 3.41 | 3.67 | 11552 | .122 | 0.003 | .002 |
| | MTG (temporal pole) | L | -51.8 | -0.13 | -19.8 | 2.66 | 3.29 | 2.81 | 5632 | .070 | 3.869 | .003 |

**Abbreviation:** ACC: anterior cingulate gyrus; Avg.: averaged; CN: caudate nucleus; H: hemisphere; IC: internal capsule; IFG: inferior frontal gyrus; IFS: inferior frontal sulcus; IOG: inferior occipital gyrus; IPL: inferior parietal lobe; L: left hemisphere; MFG: middle frontal gyrus; MOG: middle occipital gyrus; MTG: middle temporal gyrus; MTS: middle temporal sulcus; R: right hemisphere; SD: standard deviation; SFG: superior frontal gyrus; SFS: superior frontal sulcus; SMG: supramarginal gyrus; STS: superior temporal sulcus; SubACC: subgenual anterior cingulate cortex; TAL: Talairach.

**Table SR6. Clusters resulting from the searchlight RSA of emotional categories.** The resulting maps were z-transformed for each participant and, subsequently, a group-level one-sample t-test against 0 was performed (2-tailed). The clusters displayed are at uncorrected p-value = 0.005 and cluster threshold = 50 voxels.

| Location | H | TAL mean x | TAL mean y | TAL mean z | SD-x | SD-y | SD-z | Cluster size ($mm^3$) | Avg. r-value | Avg. t-value | Avg. p-value |
|---|---|---|---|---|---|---|---|---|---|---|---|
| Middle temporal gyrus | R | 61.27 | -13.3 | -17.45 | 1.91 | 2.7 | 1.86 | 3128 | 0.068 | 3.967 | 0.002 |
| Amygdala | R | 24.43 | -0.88 | -17.26 | 0.93 | 1.29 | 0.98 | 336 | -0.051 | -3.673 | 0.003 |
| Superior frontal gyrus | R | 10.16 | 40.95 | 30.24 | 1.24 | 1.56 | 1.02 | 504 | 0.071 | 3.653 | 0.003 |
| Gyrus rectus | R | 6.2 | 28.84 | -7.76 | 1.57 | 1.71 | 1.44 | 1120 | 0.054 | 3.885 | 0.002 |
| Cingulate gyrus | L | -3.19 | 18.34 | 28.05 | 1.43 | 1.06 | 1.01 | 512 | 0.064 | 3.607 | 0.004 |
| Superior frontal sulcus | L | -15.97 | 34.74 | 41.83 | 0.85 | 1.18 | 1.46 | 464 | 0.060 | 3.645 | 0.003 |
| Middle occipital gyrus | L | -28.41 | -90.07 | -4.76 | 2.74 | 3.23 | 3.35 | 5960 | 0.061 | 4.082 | 0.002 |
| Anterior insula | L | -29.79 | 24.9 | 10.1 | 2.1 | 2.02 | 1.85 | 2408 | -0.058 | -4.160 | 0.002 |
| Inferior precentral sulcus | L | -31.8 | 1.7 | 25 | 1.34 | 1.5 | 1.22 | 656 | -0.051 | -3.733 | 0.003 |
| Middle temporal sulcus | L | -41.07 | -55.09 | -2.25 | 1.74 | 1.59 | 0.92 | 736 | -0.056 | -3.856 | 0.003 |
| Middle occipital gyrus | L | -43.14 | -84.01 | -6.36 | 2.11 | 2.45 | 1.41 | 2048 | -0.051 | -3.989 | 0.002 |
| Middle temporal gyrus | L | -58.72 | -38.48 | -9.71 | 1.11 | 0.94 | 1.72 | 552 | 0.049 | 3.804 | 0.003 |
| Inferior temporal gyrus | L | -62.27 | -29.46 | -18.54 | 1.01 | 1.08 | 2.7 | 328 | -0.041 | -3.697 | 0.003 |

**Abbreviation:** Avg.: averaged; H: hemisphere; L: left hemisphere; R: right hemisphere; SD: standard deviation; TAL: Talairach.

**Table SR7. Clusters resulting from the GNB classification of emotion from functional data.** The unsmoothed functional data of each individual participant served as input for the GNB classifier, separately. Subsequently, a group-level one-sample t-test against 0 was performed (2-tailed). The reported clusters are at uncorrected p = 0.005. No cluster survived correction for multiple comparisons (cluster size corrected with Monte-Carlo simulation, alpha level = 0.05, initial p = 0.005, numbers of iterations = 5000).

| **Location** | **H** | **TAL mean x** | **TAL mean y** | **TAL mean z** | **SD-x** | **SD-y** | **SD-z** | **Cluster size ($mm^3$)** | **Avg. r-value** | **Avg. t-value** | **Avg. p-value** |
|---|---|---|---|---|---|---|---|---|---|---|---|
| IFG | R | 50.86 | 7.04 | 2.43 | 1.25 | 2.24 | 1.25 | 976 | 0.034 | 3.817 | 0.003 |
| Angular gyrus | R | 45.45 | -61.79 | 33 | 1.69 | 2.43 | 1.66 | 1632 | 0.037 | 3.807 | 0.003 |
| Fusiform gyrus | R | 20.59 | -58.14 | -12.85 | 1.71 | 1.65 | 1.44 | 1336 | 0.037 | 4.041 | 0.002 |
| ACC | R | 2.03 | 42.26 | 13.33 | 1.31 | 1.77 | 1.63 | 1072 | 0.026 | 3.772 | 0.003 |
| Angular gyrus | L | -40.2 | -67.68 | 35.68 | 2.54 | 2.01 | 1.48 | 1136 | 0.030 | 3.591 | 0.004 |
| SMG | L | -51.55 | -50.79 | 21.13 | 1.76 | 1.23 | 1.06 | 680 | 0.032 | 3.722 | 0.003 |
| MTS | L | -51.5 | -16.15 | -15.2 | 1.93 | 1.1 | 2.27 | 880 | 0.030 | 3.772 | 0.003 |
| SMG | L | -57.39 | -54.73 | 27.15 | 2.72 | 1.38 | 2.01 | 1576 | 0.035 | 3.882 | 0.003 |
| ITG | L | -58.92 | -44.43 | -14.15 | 1.43 | 3.21 | 1.22 | 824 | 0.038 | 3.619 | 0.004 |

**Abbreviation:** ACC: anterior cingulate gyrus; Avg.: averaged; H: hemisphere; IFG: inferior frontal gyrus; ITG: inferior temporal gyrus; L: left hemisphere; MTS: middle temporal sulcus; R: right hemisphere; ; SD: standard deviation; SMG: supramarginal gyrus; SubACC: subgenual anterior cingulate cortex; TAL: Talairach.

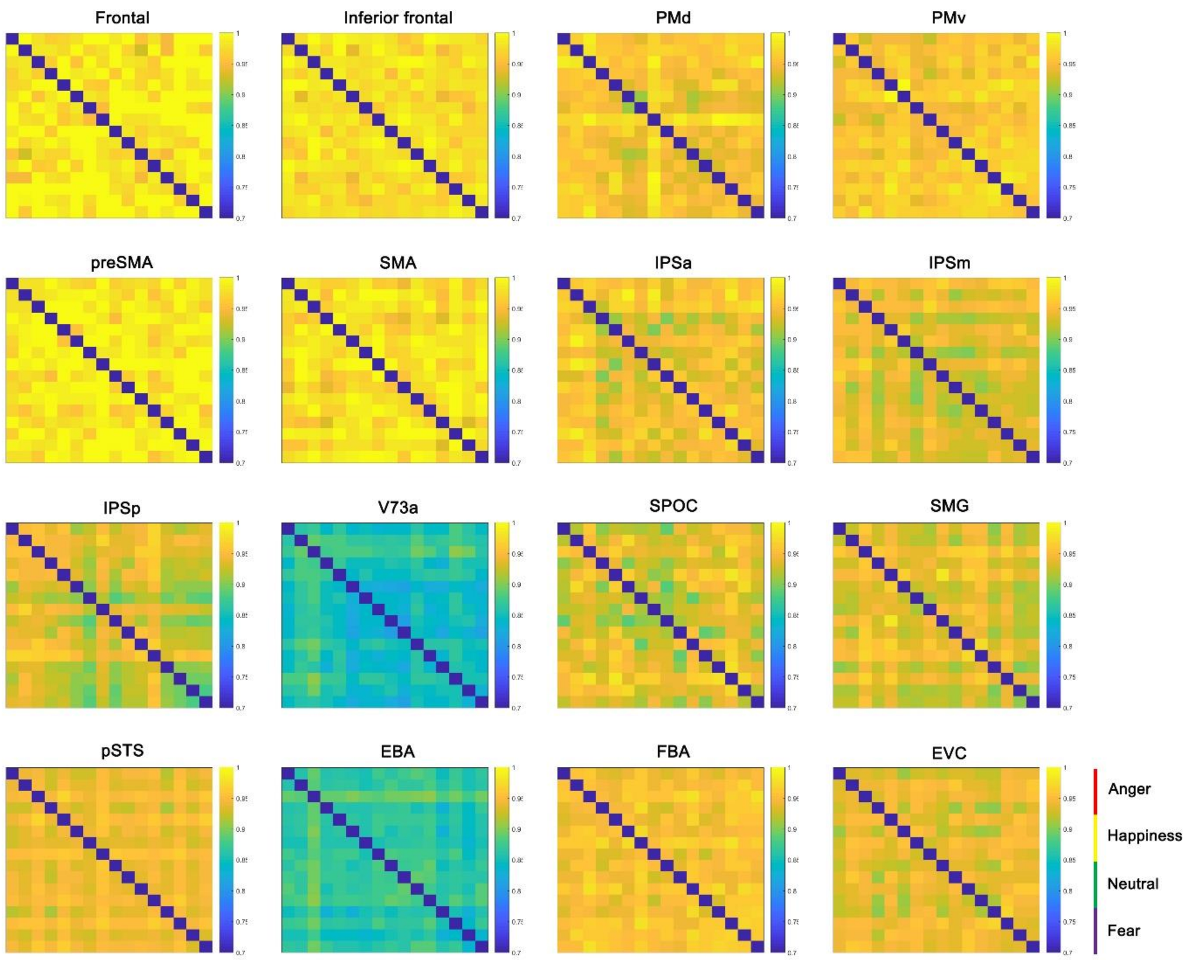


**Figure SR1. Neural representation of the affective body movements for each defined ROI.** Each matrix shows the pairwise comparisons between the neural representations of the 16 body-movement stimuli for a specific ROI. The dissimilarity measure reflects 1 - minus the sample correlation between points, with blue indicating strong similarity and yellow strong dissimilarity. Colour lines in the lower right corner indicate the organization of the RDMs with respect to the emotional category (anger: red; happiness: yellow; neutral: green; fear: purple) of the video stimuli. Order or relationships across ROIs are not assumed here. Abbreviations: EBA: extrastriate body area; FBA: fusiform body area; IFG: inferior frontal gyrus; IPS: intraparietal sulcus, p: posterior, m: middle, a: anterior; PMd: dorsal premotor cortex; PMv: ventral premotor cortex; preSMA: pre-supplementary motor area; pSTS: posterior superior temporal sulcus; SMA: supplementary motor area; SMG: supramarginal gyrus; SPOC: superior parietal occipital cortex.

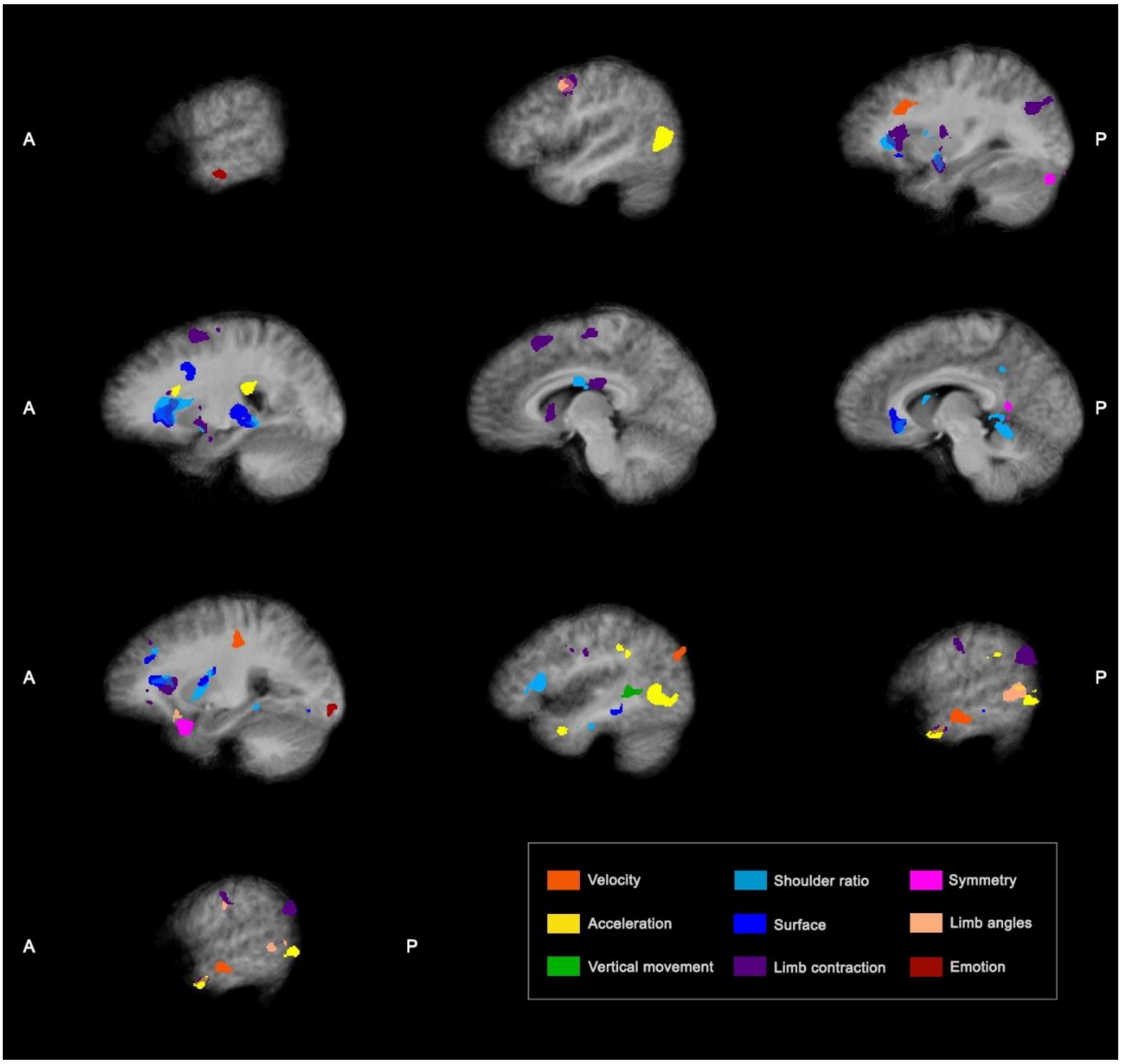


**Figure SR2. Clusters resulting from the searchlight RSA of the kinematic and postural features.** The multi-voxel fMRI dissimilarity matrices were correlated with each of the computed feature RDMs (i.e. Kinematic features: velocity, acceleration and vertical movement; Postural features: shoulder ratio, surface, limb contraction, symmetry and limb angles), respectively. The resulting maps were z-transformed for each participant. Subsequently, group-level one-sample t-tests against 0 were performed, one for each feature (2-tailed, cluster size corrected with Monte-Carlo simulation, alpha level = 0.05, initial p = 0.005, numbers of iterations = 5000). Here, both positive and negative correlated clusters are shown for each feature for a visualization of their location (see Table SR5 in Supplementary Results to see their correlation directionality).

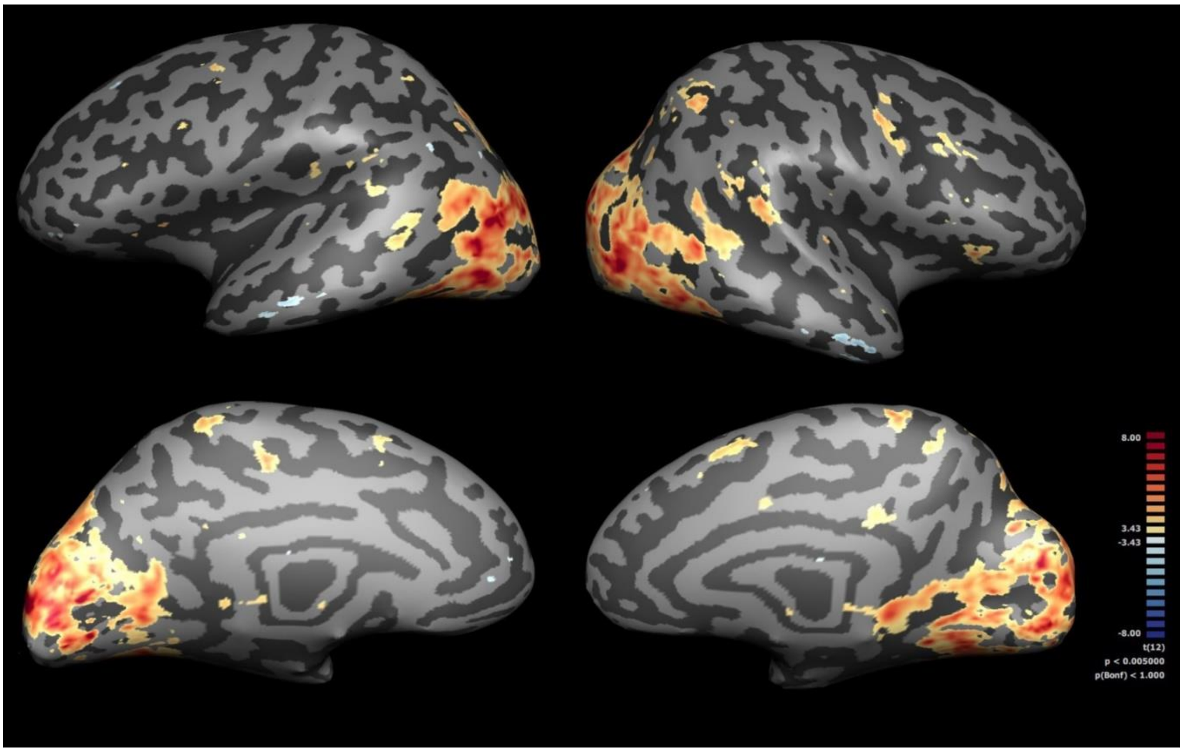


**Figure SR3. Univariate results at the group level for the body conditions.** A fixed-effects whole-brain general linear model (GLM) was fitted to the 3mm-smoothed data of the main experiment. The figure shows the univariate activation of the body conditions (i.e. body angry, body happy, body neutral and body fear) compared against baseline at uncorrected p = 0.005.
